# Multiwell-based G0-PCC assay for radiation biodosimetry

**DOI:** 10.1101/2024.05.27.596074

**Authors:** Ekaterina Royba, Igor Shuryak, Brian Ponnaiya, Mikhail Repin, Sergey Pampou, Charles Karan, Helen Turner, Guy Garty, David J. Brenner

## Abstract

In cytogenetic biodosimetry, assessing radiation exposure typically requires over 48 hours for cells to reach mitosis, significantly delaying the administration of crucial radiation countermeasures needed within the first 24 hours post-exposure. To improve medical response times, we incorporated the G0-Premature Chromosome Condensation (G0-PCC) technique with the Rapid Automated Biodosimetry Tool-II (RABiT-II), creating a faster alternative for large-scale radiation emergencies. Our findings revealed that using a lower concentration of Calyculin A (Cal A) than recommended effectively increased the yield of highly-condensed G0-PCC cells (hPCC). However, integrating recombinant CDK1/Cyclin B kinase, vital for chromosome condensation, proved challenging due to the properties of these proteins affecting interactions with cellular membranes. Interestingly, Cal A alone was capable of inducing chromosome compaction in some G0 cells even in the absence of mitotic kinases, although these chromosomes displayed atypical morphologies. This suggests that Cal A mechanism for compacting G0 chromatin may differ from condensation driven by mitotic kinases. Additionally, we observed a correlation between radiation dose and extent of hPCC chromosome fragmentation, which allowed us to automate radiation damage quantification using a Convolutional Neural Network (CNN). Our method can address the need for a same-day cytogenetic biodosimetry test in radiation emergency situations.

## Introduction

In the current global circumstances, the risk of large-scale ionizing radiation exposure due to military conflicts, terrorist activities, and accidents remains a constant concern [1]. Despite this, the general population lacks access to radiation-monitoring devices, which are typically only available to specialized personnel. This underscores the need for developing and refining biodosimetry methods to estimate radiation doses retroactively using readily accessible biological samples, such as blood. Rapid assessment of radiation doses is essential for effective emergency responses and timely medical interventions [2].

Conventional methods to access radiation damage in blood, such as the Dicentric Chromosome Assay (DCA) and the micronucleus test (MN) [3], require extensive technical work and time. They depend on cells reaching metaphase, taking over 48 hours to produce results. This delay impedes the timely administration of crucial radiation countermeasures, effective mostly within the first 24 hours post-exposure. To address these limitations, we developed a fully automated solution, the Rapid Automated Biodosimetry Tool-II (RABiT-II) [4,5], which has significantly accelerated sample processing. However, our fastest assay (the RABiT-II Dicentric Assay [6]) still requires approximately 54 hours (a total turnaround time from blood collection to result).

To further reduce the turnaround time by over 48 hours, we initiated implementation of G0-Premature Chromosome Condensation (G0-PCC) into the RABiT-II framework. G0-PCC allows to visualize chromosomal damage in non-dividing lymphocytes before mitosis, potentially enabling same-day radiation exposure assessments. There are two methods to induce G0-PCC: fusion-induced and chemically-induced [7]. Among these, fusion-induced G0-PCC is the most efficient method to condense chromosomes and is compatible with multiwell plate applications [8]. However, integrating this method into the RABiT-II may be challenging because cell fusion requires specific materials and conditions, including live, dividing hamster cells [9]. This would be difficult when needing to analyze thousands of samples within 24 hours post-exposure.

In contrast, chemically-induced G0-PCC offers a simpler way to achieve the same goal. It attempts to recreate the natural balance of phosphatases and kinases during mitosis [10] using protein phosphatase inhibitors such as Calyculin A (Cal A) and recombinant cyclin-dependent kinase 1 cyclin B (CDK1/Cyclin B) [11]. This method will allow to stockpile frozen reagents, enhancing its practicality for emergency preparedness and response. However, the chromosomes formed through chemically-induced G0-PCC do not condense as tightly as those in natural mitosis, resulting in “fuzziness” that may affect the accuracy of chromosome analysis and radiation dose reconstruction [12,13].

Recent literature has led us to hypothesize that the chemically-induced G0-PCC may not condense chromosomes as effectively as in mitosis due to the complex regulatory mechanisms of cell division and DNA supercoiling [14–16]. Research on human condensin, essential for DNA condensation [17], indicates that activating this complex in mitosis requires multiple kinases, not just CDK1/Cyclin B [18–20]. Unlike fusion-induced G0-PCC, which directly supplies resting cells with all necessary kinases, chemically-induced G0-PCC fails to introduce this diverse array of mitotic factors into cells. Additionally, the uptake of large, charged proteins like recombinant CDK1/Cyclin B (92 kDa) by G0 cells is potentially challenging, further limiting the effectiveness of premature condensation [21,22]. Considering this potential for further refinement, we prioritized chemically-induced G0-PCC with logistical considerations critical in large-scale emergencies.

An important factor to consider in integrating this method into fully automated framework is the new approach for analyzing PCC chromosomes. Previously, to analyze mitotic chromosomes, our system used custom software, which utilizes traditional image analysis like adaptive thresholding and binary object recognition. This software requires the chromosomes to be approximately straight to identify the damage using centromere and telomere spots. However, it is not as effective for identifying the less distinct chromosome patterns produced by chemically-induced G0-PCC. This limitation prompted us to start exploring more advanced deep learning techniques, such as Convolutional Neural Networks (CNN) [23–25], to improve the analysis and estimation of radiation doses in same-day test.

The use of CNN for radiation biodosimetry is gaining recognition [26–28]. CNNs are frequently employed for image analysis tasks, such as identification of objects and human faces, classification of images based on their content, and detection of diseases in medical images. Their history extends back several decades, to the work of machine learning pioneer Yann LeCun in the 1980s and 1990s [29,30]. Clearly, CNNs have strong potential to be used in radiation biodosimetry to analyze images of irradiated cells. Unlike rule-based algorithms, CNNs learn directly from the data, enabling them to recognize patterns and details that are not easily detectable by traditional methods. CNNs apply a filter to an input image to create a feature map that summarizes the presence of detected features. Importantly, CNNs do not use pre-coded filters created by humans; instead, they learn the filters during training through backpropagation and gradient descent. This significant advantage allows CNNs to automatically learn features that are specific to the task at hand, without requiring any prior knowledge or assumptions about the data.

In this study, we optimized chemically-induced G0-PCC in multiwell plates and developed a CNN-based approach for PCC image analysis and dose reconstruction (Supplementary Fig.S1). To our knowledge, this is the first attempt to combine CNNs with this type of assay for radiation biodosimetry. By training CNNs with a dataset of labeled images of both control and irradiated PCC cells, the network identified chromosomal changes induced by radiation, addressing challenges of manual analysis. These modifications significantly reduced our previous test turnaround time from 54 hours to approximately 7-9 hours, significantly improving response capability in emergency radiation exposure scenarios.

## Results

### Optimization of Protein Phosphatase Treatment

To optimize chemically-induced G0-PCC in multiwell plates, we reduced the volumes of all test components, similar to our previous approach with the DCA [6]. Initially, we used 50 nM of the phosphatase inhibitor Cal A based on prior publications (Table 1, column 3). However, observations after treatment revealed varying degrees of chromatin condensation; some cells were weakly condensed, while others were more packed (Fig. 1a). Consistent with previous reports, most of the PCC cells exhibited DNA that was not tightly packed, resulting in “fuzzy” and not fully separated chromosomes (Fig. 1a, panels i and ii). Only a few PCC cells showed completely separated chromosomes which we refer in this study as *highly-condensed PCC* (hPCC) cells (Fig. 1a, panel iv). These hPCC cells were particularly of interest because clearly separated chromosomes are easier to analyze in fully automated settings.

**Figure 1.**
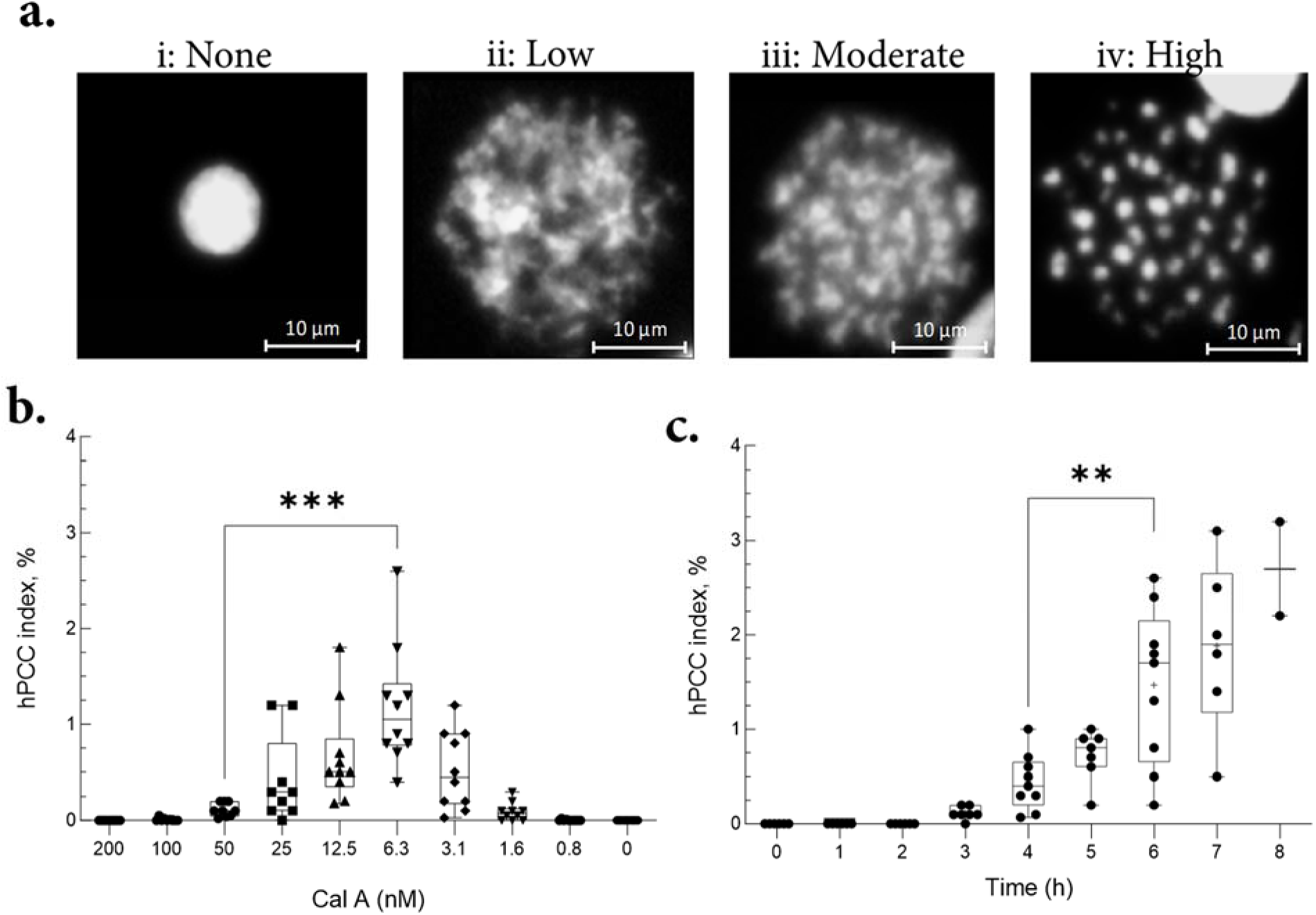
G0-Chromatin Morphologies and hPCC Index after Cal A Treatment: (a) The degrees of chromatin compaction after Cal A treatment can be categorized into four distinct categories (from left to right): (i) Intact G0 cells that show no signs of PCC and display normal, uniformly distributed chromatin typical of interphase. (ii) PCC with minimal chromatin condensation. The chromosomes are slightly more distinct than in G0 but not separated; (iii) PCC with moderate chromatin condensation. Chromatin begins to condense more significantly, showing clearer but still connected chromosomes; (iv) PCC with high chromosome condensation (hPCC). These cells display highly condensed chromosomes, with clear separation and distinct chromosomes. hPCC condensation is optimal for fully automated chromosome analysis. (b) Average hPCC index after 6 hours of treatment with various concentrations of Cal A (1-2x10^5^ cells per well). The median values from 10 independent experiments (6 donors, fresh and overnight stored blood) are shown on the bars by straight lines. The difference between hPCC index at 6.3 nM Cal A (1.18% ± 0.2 (s.e.m., n = 10)) vs. 50 nM Cal A (0.11% ± 0.02 (s.e.m., n = 9)) was statistically significant (P-value = 0.0001, unpaired t-test). (c) The effects of culture time on hPCC index at 6.3 nM Cal A. The median value from 9 independent experiments (6 donors, fresh blood only) shown by straight lines. The difference between 4 h vs. 6 h is significant (n = 9, P-value = 0.0058, unpaired t-test).

**Table 1.** Summary of Past Research On Chemically-Induced G0-PCC Assay Optimization. Various conditions tested by other research groups to tune the balance between phosphatases (Column 3) and kinases (Column 4) to ensure that cells can effectively enter premature mitosis and achieve condensation. The resulting indices of hPCC cells were generally small (Column 6), indicating potential for further optimization.

| Year | Publication | PPI | CB Kinase (if used) | Time | hPCC |
| --- | --- | --- | --- | --- | --- |
| 1999 | Kanda R. et al. [32] | OA: 200-2000 μM<br>Cal A: 50-5000 nM | | 5 h | 0.5% |
| 2000 | Prasanna P. et al. [33] | OA: 0.75 μM | 50 U | 3 h | n/d |
| 2007 | Tsuji S. et al. [21] | Cal A: 50-500 nM | 50-100 U | 6 h | 0.1% - 0.3% |
| 2014 | Pathak R. et.al. [34] | Cal A: 50 nM | 50 U | 3 h | n/d |
| 2015 | Sommer S. et al. [22] | OA: 0.75 μM<br>Cal A: 50 nM | 0-20 U | 0.45 to 3 h | 0.0% |
| 2023 | Ling G. et al. [35] | Cal A: 50 nM | 50 U | 3 to 12 h | n/d |
| 2024 | This study | Cal A: 6.3 nM |  | 4 to 6 h | 1.2% |

Abbreviations: PPI – protein phosphatase inhibitors; OA – okadaic acid; Cal A – Calyculin A; CB – recombinant CDK1/Cyclin B; U – units; hPCC – index of highly-condensed G0-PCC cells; n/d – not determined or not mentioned in the paper, but referred to as very small by the authors.

We focused on optimizing conditions to increase the number of hPCC cells per well. For this, we conducted Cal A titration tests ranging from 6400 nM to 0.8 nM. This test revealed that lymphocytes are highly sensitive to Cal A with no condensation above 100 nM (Fig. 1b and Supplementary Table S1). Interestingly, the commonly recommended concentration of 50 nM (Table 1, column 3) was suboptimal for inducing hPCC in our scaled-down tests. Instead, a much lower concentration of 6.3 nM resulted in a significantly higher hPCC index compared to 50 nM. This finding was reproducible, but only when maintained a consistent number of initial cells per well (1 to 2x10^5^ cells, see Methods). Variations in cell count per well at 6.3 nM Cal A led to non-reproducible hPCC induction, likely due to an imbalance in the ratio of Cal A molecules per lymphocyte.

We also investigated the optimal duration for 6.3 nM Cal A treatment and found that hPCC began to emerge about 3 hours after treatment initiation, peaking between 6 and 7 hours (Fig. 1c). Analyzing cells after 8 hours was challenging due to cell debris and/or reduced cell counts, aligning with previous reports that prolonged exposure to Cal A can trigger cell death [21,31].

### Optimization of Kinase Treatment

Next, we explored if adding mitotic kinase (recombinant CDK1/Cyclin B) could further improve hPCC index in our samples. Previous studies indicated that using 50 units (U) of CDK1/Cyclin B kinase could improve chromosome condensation (Table 1, column 4). However, due to the limited assay volume (300 µL) and available concentration of commercial kinase, we could only add up to 9.5 mU of kinase per well (Supplementary Methods). Since the kinase is pre-diluted in a stabilizing buffer, attempting higher concentrations would replace Cal A cell culture medium and disrupt PCC induction. Adding between 2.37 and 9.5 mU of kinase (32 or more µL per well) resulted in a non-significant decrease in PCC indices (Supplementary Fig. S2a), likely due to some cytotoxicity of kinase storage buffer components, such as Tris, EDTA, DTT, and Triton-X. Furthermore, we observed no improvement in hPCC index at lower kinase concentrations, likely due to difficulties of kinase delivery into cells that prevents to achieve mitotic threshold of kinase/cell.

To enhance kinase delivery, we developed and integrated a custom protocol into the RABiT-II framework using bacterial toxin Streptolysin O (SLO) to create pores in cell membranes, facilitating the entry of large proteins [36–38]. This method involves making cells permeable in a calcium-free environment and then reintroducing calcium to reseal the cells (Supplementary Fig. S2b-c and Supplementary Table S2). Several kinases that may play a role in facilitating mitotic chromosome condensation were added during permeabilization (CDK1/Cyclin B, CDK1/Cyclin A2, Aurora A, Aurora B, and PLK1) [39–45]. Although this treatment allowed the cells to remain viable, it did not improve the hPCC index or visibly enhance chromosome condensation.

### Morphologies of hPCC Chromosomes

During our experiments, we noticed that hPCC chromosomes, induced by Cal A without mitotic kinase, displayed unusual morphologies compared to mitotic chromosomes (Fig. 2):

- Number of Telomeres: Unlike mitotic chromosomes, which display four telomeres, each hPCC chromosome had only two, confirming the absence of DNA replication.
- Location of Telomeres: These two telomeres in hPCC were predominantly located on one side of the chromosome.
- Chromosome Sizes: hPCC chromosomes were smaller than their mitotic counterparts (e.g., hPCC chromosome 1 measured 1.9 µm across 50 hPCC cells, vs. 11.4 µm for mitotic chromosome 1 across 50 metaphase cells).
- Chromosome Lengths Uniformity: Lengths of chromosomes in hPCC were more consistent compared to the variability in mitotic chromosomes.

**Figure 2.**
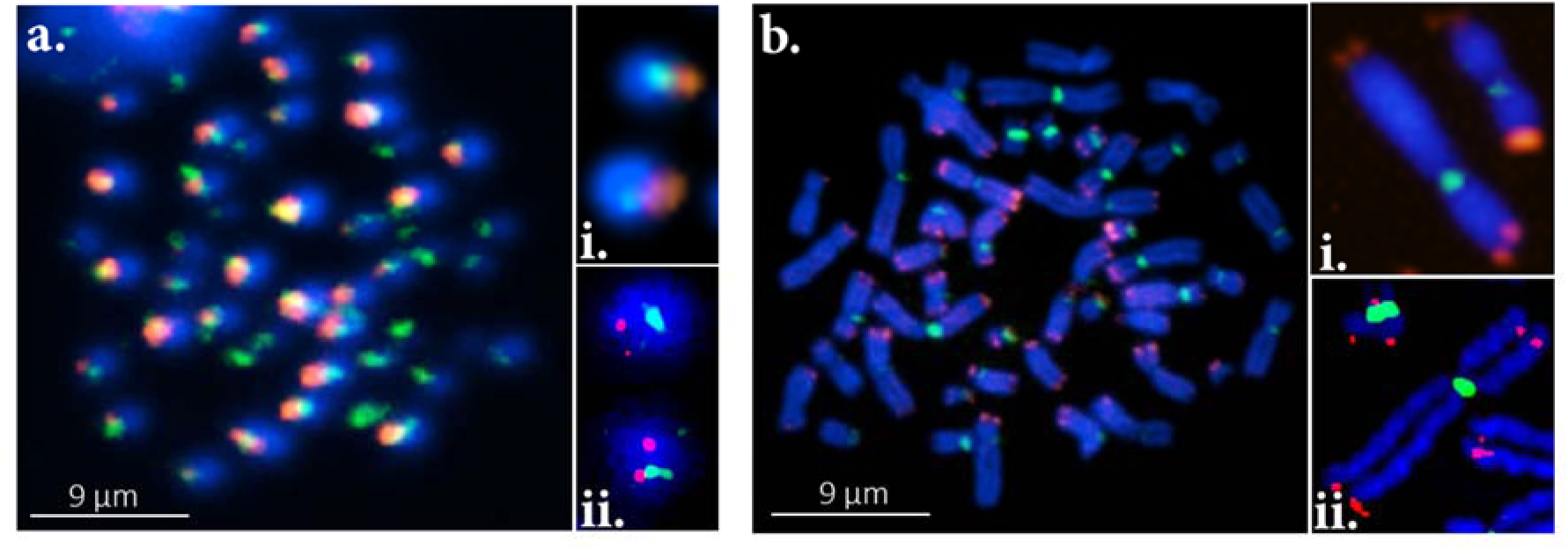
Morphologies of hPCC Chromosomes formed after treatment with Cal A without mitotic kinases: (a) hPCC chromosomes and (b) Mitotic chromosomes. Images captured at varying magnifications (20x, 40x, 63x) using BioTek and Metafer, stained with PNA FISH to highlight telomeres (red) and centromeres (green), and DNA (blue).

These unique features of hPCC chromosomes were not explored by other researchers (listed in Table 1). Thus, we continued to analyze the impact of radiation on hPCC chromosome structure and integrity. In samples exposed to low to moderate radiation doses (up to 4 Gy), identifying hPCC cells at 20x magnification by human eye was straightforward. However, at higher doses (above 6 Gy), hPCC identification by eye became difficult. Initially, it appeared there were fewer hPCC cells at higher doses. Yet, further analysis of numerous samples revealed that some cells retained distinct hPCC characteristics, such as fully separated chromosomes, though they contained more and smaller chromosomes compared to unirradiated hPCC (Fig. 3a), making them appear denser and compact (Supplementary Fig. S3), complicating identification at 20x.

**Figure 3.**
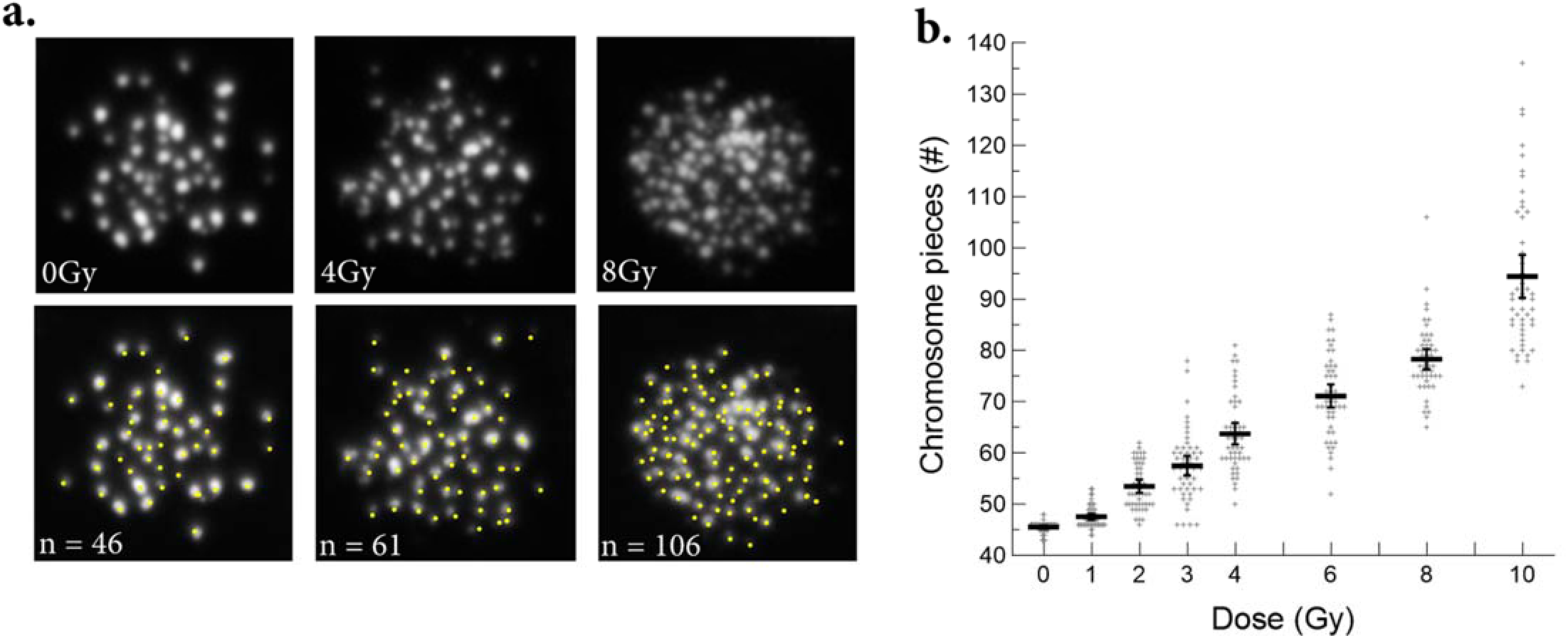
Correlation Between Radiation Dose and Chromosomal Fragmentation in hPCC. (a) Random images of hPCC from control (0 Gy) and irradiated samples (4 Gy, 8 Gy) at 20x magnification. Chromosomal pieces are marked by a scorer with yellow dots, with totals displayed below each image. (b) Fragmentation was scored in hPCC figures after exposure to 320-kVp X-rays (LET of 0.4 keV/µm). The graph shows the count of chromosomal pieces from 50 hPCC figures for each dose (Supplementary Table S3), with the average per dose represented by a straight line. Due to limitations in imaging resolution, when two pieces were close together, they were counted as one, making the counts at higher doses approximate and might not accurately reflect the true number of broken chromosome pieces per hPCC.

To assess whether irradiated hPCC cells indeed exhibited increased chromosomal fragmentation, we manually counted chromosome pieces per hPCC across doses from 0 to 10 Gy (Supplementary Fig. S4). This analysis indicated a potential correlation between radiation dose and the number of chromosomal pieces per hPCC (Fig. 3b and Supplementary Table S2).

### Dose Reconstruction

To analyze hPCC chromosomes, we initially tried our existing RABiT-II chromosome identification software, but encountered several challenges when adapting the classifier for hPCC chromosomes:

- Limited hPCC index: The software requires a larger dataset for optimal performance. However, hPCC index is lower than mitotic index.
- Sample Preparation Difficulty: Extracting hPCC chromosomes from cell nuclei and preparing them for analysis proved challenging.
- Radiation Damage: Exposure to radiation often resulted in broken chromosomes in hPCC cells, complicating visualization of centromeres with PNA FISH.
- Unique Features: hPCC chromosomes exhibit distinct shapes and features not found in mitotic samples.

Given these obstacles, we decided to modify our chromosome analysis approach by employing a deep learning model tailored to handle these unique characteristics of hPCC chromosomes. After CNN training, we conducted a preliminary double-blind dose-reconstruction test. Blood samples from a healthy 35-year-old male volunteer were irradiated with three different doses (labeled X, Y, and Z) and processed by different researchers (Researcher 1: irradiation; Researcher 2: sample processing and hPCC image collection; Researcher 3: sample analysis) to ensure blinding. Approximately 200 hPCC images were collected for each sample. Researcher 3 used predicted class probabilities for each image to generate dose reconstructions, which were then decoded by Researcher 1. Preliminary test data is summarized in Table 2, showing predicted values as mean ± standard deviation. The results demonstrated that the median and mean dose reconstruction values were reasonably close to the actual doses, encouraging further refinement of the model.

**Table 2.** Independent Validation Results for the original CNN model version with 4 dose classes (non-reacted, 0 Gy, 3 Gy and 8 Gy) on three data sets (labeled X, Y and Z).

| Data set | Actual Dose (Gy) | Number of images | Summary statistics for reconstructed dose |  |  |  |  |  |  |
| --- | --- | --- | --- | --- | --- | --- | --- | --- | --- |
|  |  |  | minimum | 25th percentile | 50th percentile (median) | 75th percentile | maximum | mean | Standard deviation |
| Y | **0** | 183 | 0.18 | 0.45 | **1.01** | 1.79 | 6.31 | 1.46 | 1.33 |
| Z | **2.5** | 239 | 0.36 | 0.87 | **1.75** | 3.58 | 7.33 | 2.29 | 1.68 |
| X | **6** | 207 | 1.91 | 5.26 | **6.37** | 7.06 | 7.80 | 6.01 | 1.33 |

Next, the CNN model was expanded to include 10 dose classes (non-reacted, 0, 1, 2, 2.5, 3, 4, 6, 8, and 10 Gy) and enhanced with quantile regression for correction. This refined architecture includes:

- Five convolutional layers with varying dimensions and Leaky ReLU activation functions.
- Two fully connected layers with Leaky ReLU activation.
- Dropout layers for regularization and noise augmentation for robustness.
- Custom image augmentation classes, ’AddGaussianNoise’ and ’BlackoutCircle’, to introduce Gaussian noise and simulate blackout circles on images, respectively.
- A ’cutmix_data’ function to apply CutMix augmentation, enhancing data diversity.

Examples of augmented images generated for CNN training are shown in Supplementary Fig. S5. The total expanded dataset comprised 6,030 images. We randomly allocated 70% of these images to the training set, with the remaining 30% designated for the testing set. Fig. 4a displays random examples of images from the 0, 6, and 10 Gy classes. Upon visual examination, there appears to be a trend of increasing fragmentation and “fuzziness” of the chromosomes as the radiation dose increases. However, it is important to note that this trend is not consistently discernible across all individual images.

**Figure 4.**
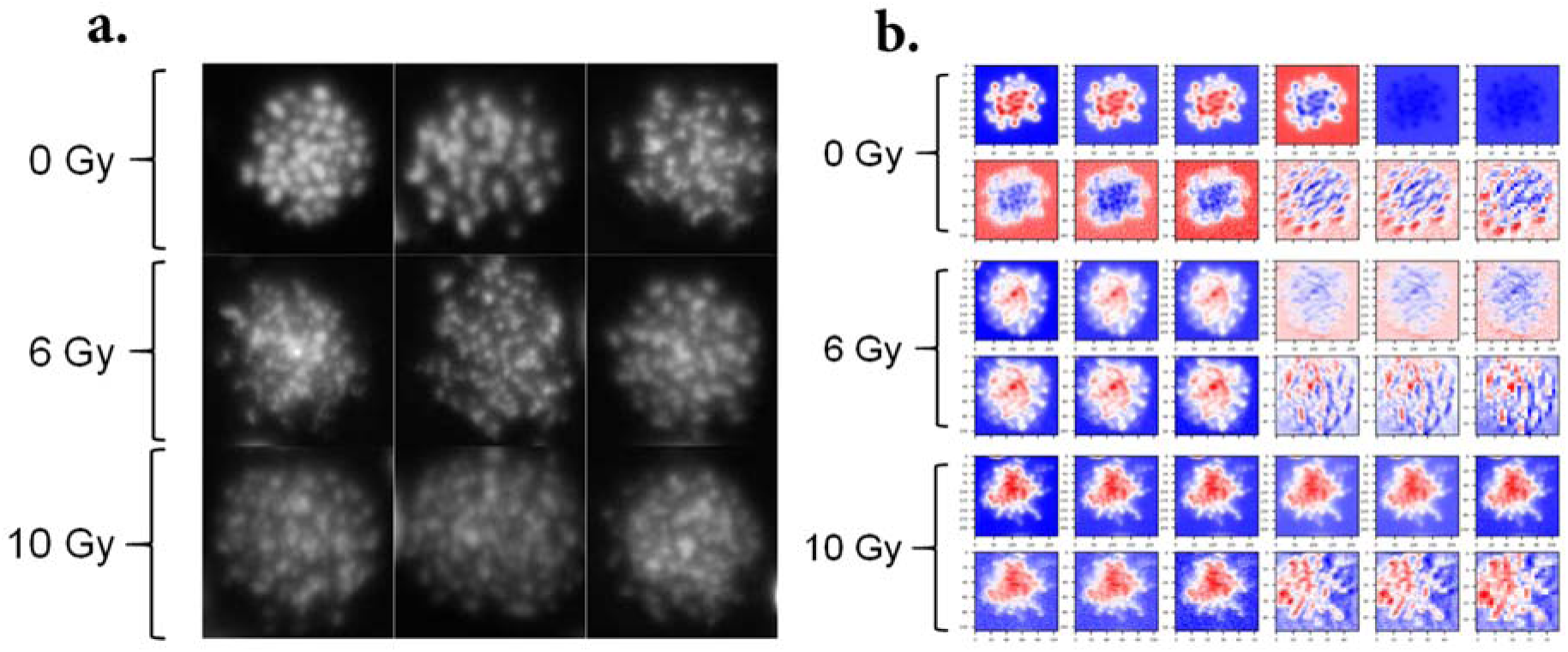
Chromosomal Variations and CNN Analysis Across Radiation Doses: (a) Three random images from the 0, 6, and 10 Gy classes are shown as examples. (b) CNN feature extraction visualization. A sequence of heatmaps from 12 modules of the CNN to illustrate the progression of feature extraction and transformation across the layers of the network. Three example images randomly selected from the 0, 6 and 10 Gy classes were used as inputs for these heatmaps. Details of the interpretation are described in the main text.

To illustrate what features the CNN model focuses on when making predictions, we generated a sequence of heatmaps from 12 modules of the CNN to illustrate the progression of feature extraction and transformation across the layers of the network (Fig. 4b). Each module in a CNN typically represents a different level of abstraction, so the heatmaps show how the network’s focus changes as it moves deeper into the model. Each heatmap is color-coded, with blue indicating lower activations and red indicating higher activations. This means that the model pays less attention to the blue regions and more attention to the red regions of the image when making its prediction. Essentially, the red areas in the heatmap are where the model sees the most evidence for the class it has predicted, while the blue areas contribute less to the model’s decision. This visualization is a useful tool for understanding what the model is “looking at” and how it is making its decisions, thereby making our models more interpretable and transparent. Although in this case intuitive interpretation of the activation maps is not straightforward, it appears that there are substantial differences in the map patterns for the 3 selected images, especially for the last 3 of the selected modules.

The performance of the refined CNN model with quantile regression correction on the expanded 10-class data set is summarized numerically in Table 3 and graphically in Fig. 5. These results support the findings of the original blinded validations described above, and suggest that the median reconstructed doses for each class are reasonably close to the actual dose values. In a practical biodosimetry situation, hundreds of PCC images from a single individual will be collected.

**Figure 5.**
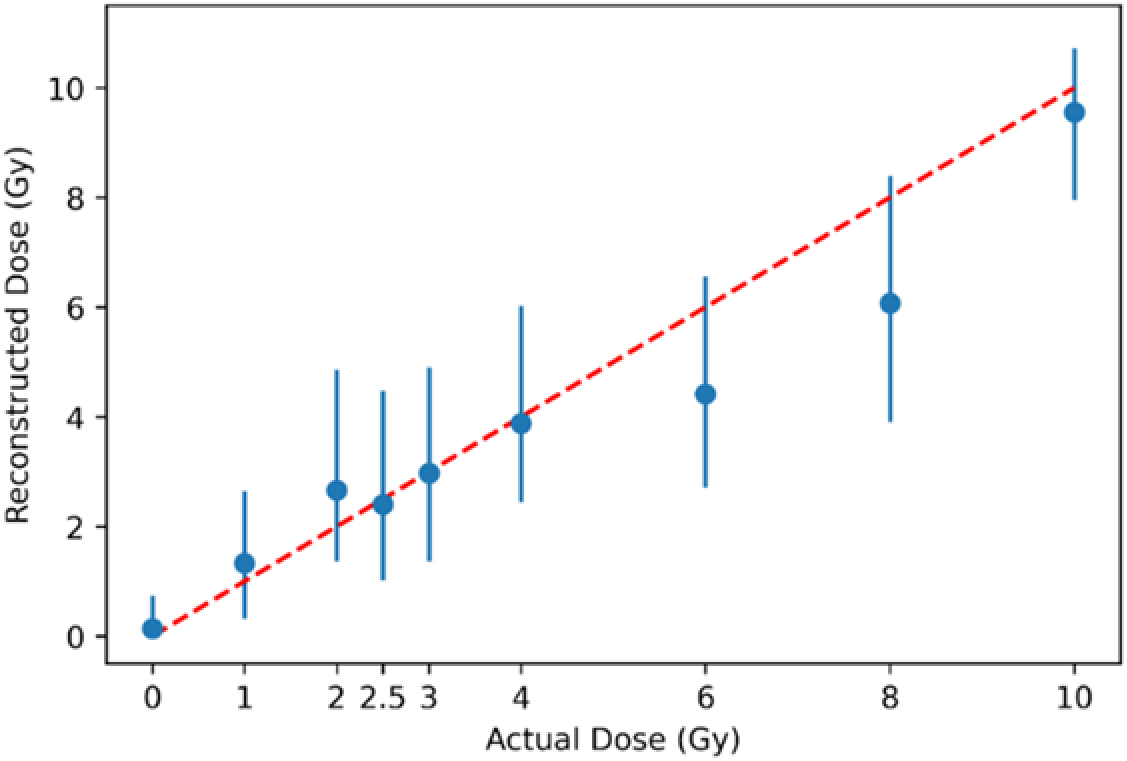
Dose Reconstruction. Visual comparison of actual and reconstructed doses generated by the refined CNN model with quantile regression correction on the testing portion of the expanded 10-class data set. Blue circles are medians and error bars represent the interquartile range (25th to 75th percentiles). The red dashed line represents theoretical 1:1 correspondence.

**Table 3.** Reconstructed Doses Statistics. Summary statistics for reconstructed doses generated by the refined CNN model with quantile regression correction on the testing portion of the expanded 10-class data set.

| Actual Dose (Gy) | Number of images in testing set | Summary statistics for reconstructed dose on the testing data |  |  |  |  |
| --- | --- | --- | --- | --- | --- | --- |
|  |  | minimum | 25th percentile | 50th percentile (median) | 75th percentile | maximum |
| 0.0 | 527 | 0.00 | 0.00 | **0.14** | 0.74 | 7.94 |
| 1.0 | 60 | 0.06 | 0.32 | **1.33** | 2.65 | 8.94 |
| 2.0 | 60 | 0.00 | 1.37 | **2.66** | 4.86 | 10.22 |
| 2.5 | 79 | 0.00 | 1.02 | **2.40** | 4.48 | 8.34 |
| 3.0 | 356 | 0.01 | 1.37 | **2.97** | 4.90 | 10.69 |
| 4.0 | 60 | 0.39 | 2.45 | **3.88** | 6.03 | 11.06 |
| 6.0 | 129 | 0.02 | 2.72 | **4.42** | 6.57 | 10.84 |
| 8.0 | 360 | 0.06 | 3.90 | **6.07** | 8.40 | 10.67 |
| 10.0 | 60 | 4.52 | 7.95 | **9.55** | 10.72 | 11.89 |

## Discussion

The RABiT-II DCA is an automated cytogenetic test developed to quickly assess hundreds of blood samples for signs of radiation damage [46]. Typically, the turnaround time of this test is about 54 hours, primarily because it takes over 48 hours for cells to reach mitosis and display chromosomal damage. To reduce this time, we integrated chemically-induced G0-PCC into a high-throughput format, aiming to enable same-day testing for rapid deployment in large-scale radiation emergencies. Optimizing PCC induction in non-dividing (G0 phase) cells requires reducing phosphatase activity and enhancing kinase activity to activate the condensin complex for mitotic-like chromosome condensation.

Our findings indicate that G0 lymphocytes are highly sensitive to even minor variations in phosphatase inhibitor concentrations. Adjusting Cal A to 6.3 nM, instead of the recommended 50 nM, significantly improved hPCC yield in a 300 µL volume. This improvement was reproducible and depended on maintaining a consistent cell concentration per well. Variations in cell counts at 6.3 nM Cal A led to inconsistent hPCC indices, highlighting the need for a balanced Cal A-to-lymphocyte ratio for consistent results. This may require tailored optimization across different laboratories.

Reproducibility tests with 6.3 nM Cal A revealed that: 1) Fresh blood consistently yielded higher hPCC indices than the same blood stored overnight at room temperature; 2) The response to Cal A varied among donors, with younger individuals generally exhibiting higher hPCC indices. Our findings also underline the importance of identifying the optimal timing for peak hPCC condensation and the potential risks of cell death from prolonged Cal A exposure. These insights are crucial for fine-tuning the G0-PCC method in practical settings, ensuring the RABiT-II system delivers timely and accurate results in radiological emergencies.

Condensin’s crucial role in chromosome condensation during mitosis is well-documented [47–49]. Recent studies have indicated that this complex requires various mitotic kinases for full activation [19]. Considering this, beyond the standard CDK1/Cyclin B, we explored additional kinases that regulate condensin at the onset of mitosis [39–45, 50]. However, introducing these kinases into the RABiT-II assay presented several challenges:

- Threshold Concentration: Achieving the necessary kinase concentration in each well without diluting essential culture medium components with chemicals from the kinase storage buffer, such as DTT and Triton X-100, was problematic.
- Cellular Transport: Another issue was ensuring kinases could penetrate the outer cell membrane and reach the nucleus in concentrations sufficient to induce mitosis [51]. It is unclear whether specific membrane transporters for these recombinant kinases exist. Fusion-induced G0-PCC directly introduces essential mitotic kinases into the nuclei of G0 cells during fusion, improving chromosome morphologies. However, the effectiveness of kinase uptake in chemically-induced G0-PCC is uncertain [22], especially given the large molecular weight of recombinant cyclins (91 kDa for CDK1/Cyclin B and 135 kDa for CDK1/Cyclin A2).

To address these issues, we utilized the bacterial toxin SLO, which binds to cholesterol in plasma membranes and forms pores up to 35 nm in diameter. This method allows large molecules (up to 150 kDa) to enter cells, and the pores can subsequently be resealed, preserving cell viability [36]. We tested this approach to enhance the entry of large protein kinases into lymphocytes. Despite using SLO, we observed no increase in hPCC indices. Efforts to monitor the movement of recombinant kinases during SLO-mediated permeabilization was unsuccessful. The commercially available kinases are offered at very low concentrations (less than 1 mg/mL), which negatively affected their labeling with a fluorescent tag needed for tracking. The failure to achieve further condensation may be due to insufficient kinase concentrations per cell, suboptimal kinase mix selection, or the kinases inability to traverse lymphocyte membranes even with SLO. Therefore, the underlying reasons for the lack of further condensation remain unclear and require further clarification.

Without kinases, unirradiated hPCC chromosomes typically exhibited 46 distinct segments, corresponding to the human chromosome count. Each segment retained essential structural features, such as a centromere and telomeres. However, hPCC chromosomes displayed notable differences in appearance compared to typical mitotic or fusion-induced G0-PCC chromosomes. This unusual morphology could be linked to the effects of Cal A on resting cells, which lack the mitotic kinases necessary for normal chromosomal condensation and nuclear envelope breakdown. Specifically, the absence of CDK1/Cyclin B, crucial for nuclear envelope disassembly and activation of the condensin complex to supercoil chromatin into “rod”-shaped chromosomes, might explain these observations [52,53]. The nuclear envelope in hPCC cells might remain at least partially intact. This could cause chromosomes to stay anchored to the envelope by their telomeres, maintaining their spatial orientation within “chromosome territories” [54,55]. This aligns with models of chromosome-telomere positioning seen in some mammalian cells [56] and could explain why hPCC telomeres were predominantly positioned on one side, and why most of hPCC cells were resistant to chromosome isolation from nuclei for automated analysis.

These findings raise several questions: If all lymphocytes were in a non-dividing (G0) state prior to Cal A addition, why did some hPCC cells exhibit tightly packed chromosomes while most showed only weak to moderate condensation? Additionally, why was the proportion of these hPCC cells relatively consistent across different donors we tested (about 1%)? Could it be that these cells were not truly in G0 but were instead cycling, equipped with internal kinases that facilitated chromosome condensation? And if so, why do these hPCC chromosomes appear atypical compared to those in cycling cells? These discrepancies require further investigation into the mechanisms behind unusual hPCC chromosome morphology.

Following irradiation, we noted a significant increase in chromosome fragmentation within hPCC figures, showing a dose-dependent correlation. Here we define fragmentation as the immediate breaking of chromosomes into smaller pieces observed within hPCC figures. This likely results from the acute impact of radiation, leading to structural breakdown and visible breaks that may not undergo normal repair processes. This contrasts with the acentric pieces and rings typically seen in 48-hour PCC tests, which result from inefficient DNA repair after irradiation. The hPCC fragmentation pattern resembled two scenarios: 1) S-phase PCC, which occurs when Cal A is added during DNA replication [57]; or 2) Chromosome Fragmentation (C-Frag), a type of cell death during mitosis where chromosomes break down under various types of cellular stress [58]. This suggests that radiation may compromise the structural integrity of hPCC chromosomes, though the exact mechanistic nature behind this fragmentation remain unclear.

Our preliminary flow cytometry data (Supplementary Fig. S6) revealed that even mild exposure to Cal A rapidly led to progressive changes in nucleosome state. When combined with radiation, these changes were accompanied by the accumulation of unrepaired DNA damage, indicated by H2AX foci. Interestingly, while cells irradiated but not treated with Cal A managed to repair such damage, those treated with Cal A did not, suggesting that Cal A may induce a spatial compression of nucleosomes. This compression could mechanically restrict DNA repair proteins from accessing and repairing newly damaged DNA sites. These observations indicate that inhibiting phosphatase activity with Cal A triggers a rapid contraction of the chromatin, potentially driven by changes in the phosphorylation states of histones. This effect is distinct from traditional chromosome condensation, mediated by condensins [59,60] activated by mitotic kinase/s. Thus, the chromatin structure induced by Cal A in hPCC may represent a form of “spatial compaction” rather than “condensation” achieved through DNA twisting and supercoiling.

To effectively analyze hPCC chromosome fragmentation, we developed a novel approach to analyze hPCC images for biodosimetry, based on a customized CNN model. This model generates a probability distribution across different image classes (representing various doses), which is converted into a continuous reconstructed dose output. The model employs multiple layers, each representing a different level of data abstraction. Visualization of these layers is achieved through heatmaps that are color-coded – blue for lower activations and red for higher activations (Fig. 4b). This color coding helped to illustrate how our model shifts its focus deeper into the network, paying more attention to hPCC areas indicated by red, which represent regions where the model found the most evidence for its predictions. Conversely, blue areas indicated lesser focus, contributing minimally to the decision-making process. These heatmaps serve as a valuable tool for understanding the CNN analytical focus and decision-making process for dose reconstruction.

Correction by quantile regression helps alleviate asymmetric error distributions. Training CNN with a diverse range of hPCC images enabled the system to analyze the complex patterns of hPCC fragmentation that our conventional RABiT-II chromosome analysis software was not able to match. Initial blind tests of this system showed reasonable performance in initial blinded tests (Table 2) and testing data from a larger dataset (Table 3 and Fig. 5). We believe this demonstrates a proof of principle for using deep learning to enhance the potential of the chemically-induced G0-PCC assay in radiation biodosimetry.

Some limitations of this study include the manual selection and cropping of hPCC images for analysis. As described, we were careful to avoid biasing the results by selecting images according to pre-determined criteria. In future developments, automated image selection techniques are planned to improve throughput capabilities and gather larger data sets. Strengths of the study include the relatively small size of the customized CNN model (described in more detail in Supplementary Methods), which allows model training on readily-available machines and rapid generation of predictions for multiple images from the trained model.

Further refinement is needed in several critical aspects of our assay. We plan to continue optimizing the delivery and functional effectiveness of kinases within the assay. Additional testing is necessary to better understand the unusual condensation and post-irradiation fragmentation patterns observed in hPCC cells. Although we have successfully reduced the total assay turnaround time from 54 hours to approximately 7-9 hours, our goal is to further decrease this duration to better suit rapid response scenarios in radiological emergencies. Moreover, we intend to improve the precision of our dose reconstruction by categorizing radiation exposure into distinct bins – unexposed (0 Gy), mild (1-2 Gy), moderate (2-4 Gy), severe (4-6 Gy), very severe (6-8 Gy), and lethal (>8 Gy) – to enhance the accuracy and relevance of our high-throughput test.

## Methods

All reagents and plasticware, unless specified otherwise, were sourced from Thermo Fisher Scientific^TM^ (Waltham, MA).

### Blood Samples

The Institutional Review Board (IRB) of Columbia University approved this study under protocol # AAAF2671 and all experiments were performed in accordance with these guidelines and regulations. All participants were healthy volunteers aged between 27 and 63 years, with no recent exposure to ionizing radiation or clastogenic agents. After giving informed consent, peripheral blood samples were collected by a certified nurse through venipuncture into vacutainers containing sodium heparin anticoagulant (Becton Dickinson, NJ). Peripheral blood mononuclear cells (PBMCs) were isolated using a density gradient (Histopaque®-1077, density: 1.077 g/mL, Sigma-Aldrich, MO), according to the provided user manual instructions. After isolation, PBMCs were washed twice with HBSS buffer (without calcium and magnesium) and resuspended in HBSS to a final concentration of 2-4x10^6^ cells/mL prior to irradiation.

### Irradiation

Cell suspensions were irradiated in 2D Matrix Storage Tubes using an X-RAD 320 (Precision X-Ray, North Branford, CT), positioned 40 cm from the X-ray target. The settings were adjusted to 320-kVp, 12.5 mA, with a 2mm Al filter, delivering radiation at a rate of 4 Gy/min. The dose and dose rate were determined using a Radcal 10x6-6 (Moravia, CA) cylindrical ion chamber and verified with a built-in parallel plate transmission chamber integrated into the filter assembly. Dose uniformity was assessed using EBT3 radiochromic film (Ashland Advanced Materials, Bridgewater, NJ). Calibration of the X-RAD 320 built-in transmission chamber was also performed using a Radcal ion chamber. Control samples were sham irradiated, receiving a 0 Gy dose.

### Reagent Reconstitution

Calyculin A (Cal A) from *Discodermia calyx* (10 µg, MW = 1009.17 g/mol, Sigma-Aldrich, MO), was dissolved in 99 µl of dimethyl sulfoxide (DMSO) to achieve a concentration of 100 µM and stored at -20°C for no longer than a month. Our trials with alternative solvents, including ethanol, a 1:1 mix of ethanol and water, and a 1:1 mix of DMSO and water, showed that DMSO provided superior solubility and absorption for Cal A. Consequently, all study results are based on Cal A dissolved in DMSO.

Phytohemagglutinin-M (PHA-M) from *Phaseolus vulgaris* (20 mg, Sigma-Aldrich, MO) was diluted in 2 ml of distilled water to a working concentration of 10 mg/ml and stored at -20°C for up to two months.

### Multiwell-Based G0-PCC Assay

We adapted the chemically-induced G0-PCC assay for a multiwell format, incorporating several modifications from the RABiT-II DCA method [6]. Briefly, 50 µl cell aliquots were combined with 250 µl of pre-warmed MarrowMAX™ Bone Marrow Medium, containing 20% heat-inactivated Fetal Bovine Serum (Gibco™), 75 µg/ml PHA-M, and 6.3 nM Cal A, resulting in approximately 1 to 2x10^5^ lymphocytes per well. The cells were incubated at 37°C in plastic 96-well plates (Corning®) for 6 hours, then washed once in 0.075M KCl, subjected to swelling in KCl for 6 minutes, and fixed using a 3:1 methanol:acetic acid mixture. After fixation, the cells were washed once in a solution of 80% acetic acid and 20% water, then immersed for 2 minutes in the same solution before transfer to glass bottom 96-well plates (DotScientific, MI). Cells were stained with 4’,6-diamidino-2-phenylindole (DAPI) dissolved in PBS, and imaging was performed using a BioTek Cytation Cell Imaging Multi-Mode Reader (Agilent Technologies Inc., CA) at 20x magnification.

### Highly-condensed PCC Index and Statistical Analysis

During the optimization of the assay, the proportion of hPCC cells per well (sample) was calculated using the following formula adapted from another study [33]:

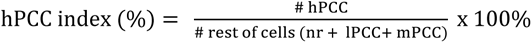

where:

- hPCC is highly-condensed PCCs,
- nr is non-reacted cells,
- lPCC is low-condensed PCCs,
- mPCC is moderately condensed PCCs per well.

All experiments included a minimum of three biological replicates. Statistical significance was assessed using an unpaired t-test, with analyses performed in GraphPad Prism v9.3.1 software.

### Selection Criteria for CNN Libraries

Selection of hPCC cells was based on distinct features: (i) clearly separated chromosomes (46 chromosomes in controls); (ii) average PCC figure diameters of over 25 µm (across 50 cells); and (iii) all chromosomes within a PCC figure measuring from 1.3 to 3 µm long (across 50 cells). To minimize bias during the accumulation of the CNN hPCC image libraries, we selected hPCC cells from different experiments based on only two main criteria: first, the chromosomes must be completely separated so that each chromosomal fragment could be manually counted if necessary; second, each PCC figure must be surrounded by a black background, free from any other cells, debris, or artifacts.

### Deep Learning Approach for Image Analysis and Dose Reconstruction

CNNs are feed-forward neural networks that refine feature engineering through filters – small matrices of numbers (e.g., 5x5) that traverse the image pixel by pixel. The values in the filter matrix are multiplied by the corresponding pixel values in the image, and the resulting products are summed to produce a single output value. The output values form a new matrix, which represents a transformed version of the original image. After the convolution step, a non-linear activation function is applied to the output matrix. This helps to introduce non-linearity into the model, which allows it to learn more complex patterns in the data. Next, a pooling layer is applied to the output of the activation function. This reduces the dimensionality of the output matrix by taking the maximum (or average) value of a small subset of the matrix. This helps to reduce the number of parameters in the model, making it more efficient. Finally, the output of the pooling layer is flattened and fed into a fully connected neural network. This part of the model is similar to a traditional neural network, and is responsible for making the final predictions based on the features learned in the convolutional and pooling layers (Supplementary Fig. S1).

During training, the CNN processes a set of labeled images, categorized into different classes, starting with randomly initialized filter values. Through forward propagation and backpropagation, the network adjusts its filters to minimize the loss function—the discrepancy between predicted outputs and actual labels—using optimization algorithms like gradient descent,

Our customized approach for image analysis and dose reconstruction is outlined in the Supplementary Methods. Key steps include:

1. *Develop a customized convolutional neural network (CNN) model to classify PCC images into dose classes*, and train this model on the training data set (randomly selected 70% of available images in each class). The output of this procedure for each image is a predicted probability distribution over all classes. Initially, there were 4 classes: non-reacted (representing a failed PCC reaction), 0 Gy, 3 Gy and 8 Gy. During subsequent model development, the number of classes was increased to 10: non-reacted, 0, 1, 2, 2.5, 3, 4, 6, 8, and 10 Gy. The CNN architecture was manually optimized in terms of number of convolutional layers, total number of trainable parameters, and dropout and Gaussian noise layers to reduce overfitting. The goal was to produce a small and quickly-implementable model with decent performance. Several image augmentation techniques were implemented during CNN training to improve performance and reduce overfitting.
2. *Convert the classification output (class probabilities) into a continuous dose reconstruction for each image*. Since the actual dose (*D_i_*) corresponding to each class (*i*) is known, the reconstructed dose (*D_R_*) can be calculated as the sum of the products of class probabilities (*P_i_*) and the actual doses (*D_i_*) over all classes (*i*=1..*N*), as follows: *D*_R_ = 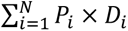. In this calculation *D_i_* for the non-reacted class was assumed to be zero.
3. *Correct the reconstructed doses using quantile regression*. The motivation for this correction procedure was that the error distributions for the reconstructed doses (*D_R_*) were not symmetrical, especially for the low and high extremes of the studied dose range. For the lowest dose class (0 Gy) the CNN could only make errors in the “positive” direction, where the probability distribution for a given image included some non-zero values for higher dose classes. For the highest dose class of 10 Gy the situation was reversed, with all errors being in the “negative” direction with some non-zero probabilities for lower dose classes. Consequently, the median *D_R_*values for low doses tended to be overestimated, whereas for high doses they were underestimated. We used quantile regression to correct for these systematic deviations as follows: A polynomial function of the *D_R_* values generated by the CNN on training data was used to predict the corresponding actual dose (*D_i_*) values, using quantile regression for the median. The best-fit parameters of the regression were recorded for use later. This correction was implemented only in the second version of the model with 10 dose classes.
4. *Generate a reconstructed dose for each image in the testing data set* using the trained CNN model to make class predictions and correcting these predictions with the quantile regression model with previously recorded parameters.
5. *Generate a reconstructed dose for an independent data set from an individual exposed to an unknown radiation dose.* In a realistic biodosimetry scenario, several hundred images from an individual will be used. The final dose reconstruction for the individual will be the median of reconstructed doses over all images from that individual.

As mentioned above, we initially used 4 image classes (non-reacted, 0 Gy, 3 Gy and 8 Gy) to train the model. There were 1000 images in each class, with 70% in each class assigned to training data and the remainder to testing data. The trained model was evaluated on the testing data. Importantly, it was also validated on three independent data sets as follows: (1) Each of these data sets (labeled X, Y and Z) was generated by irradiating blood samples from a 35-year-old male donor with a different radiation dose. (2) The PCC assay was performed and images were generated from each data set. (3) The images were from each data set were imported into the CNN model to generate dose reconstructions. In this procedure, the researcher performing the CNN-based dose reconstructions was blinded to the actual doses for data sets X, Y and Z. Following this independent validation, which performed relatively well as described in the Results section, the number of dose classes was increased to 10 (non-reacted, 0, 1, 2, 2.5, 3, 4, 6, 8, and 10 Gy) and the number of images was also increased. Subsequent rounds of model training and optimization were performed.

## Acknowledgements

The authors would like to thank Maria Taveras (Center for Radiological Research), the nurses of the Dept. of Radiation Oncology, Dr. Norman Kleiman and Lauren Stiene (Kleiman Laboratory, Columbia University) for their help with blood collection. We also grateful to Michael Kissner and Rosemary (Rose) Gordon-Schneider from the Columbia Stem Cell Initiative Flow Cytometry Core for their guidance in planning the flow cytometry experiments. This research received support from the grant U19-AI067773 (PI Dr. David J. Brenner), awarded by the National Institute of Allergy and Infectious Diseases, the National Institutes of Health.

## Author Contributions

BP and DJB conceived the idea. ER, BP, GG, MR, SP, and CK performed the experiments. IS developed the CNN model and performed statistical analyses. ER and IS wrote the main manuscript text and prepared the figures. All authors reviewed the manuscript.

## Data Availability

All data generated or analyzed during this study are included in this published article (and its Supplementary Information files).

## References

1. Shuryak, I. et al. A machine learning method for improving the accuracy of radiation biodosimetry by combining data from the dicentric chromosomes and micronucleus assays. Sci. Rep. 12, 21077 (2022).

2. Blakely, W. F., Prasanna, P. G., Grace, M. B. & Miller, A. C. Radiation exposure assessment using cytological and molecular biomarkers. Radiat. Prot. Dosimetry 97, 17– 23 (2001).

3. Testa, A., Palma, V. & Patrono, C. Dicentric Chromosome Assay (DCA) and Cytokinesis-Block Micronucleus (CBMN) Assay in the Field of Biological Dosimetry. in 105–119 (2019).

4. Garty, G. et al. The Decade of the RABiT-II (2005-15). Radiat. Prot. Dosimetry 172, 201– 206 (2016).

5. Repin, M., Pampou, S., Karan, C., Brenner, D. J. & Garty, G. RABiT-II: Implementation of a High-Throughput Micronucleus Biodosimetry Assay on Commercial Biotech Robotic Systems. Radiat. Res. 187, 502–508 (2017).

6. Royba, E. et al. RABiT-II-DCA: A Fully-automated Dicentric Chromosome Assay in Multiwell Plates. Radiat. Res. 192, 311 (2019).

7. Ravi, M., Nivedita, K. & Pai, G. M. Chromatin condensation dynamics and implications of induced premature chromosome condensation. Biochimie 95, 124–133 (2013).

8. Pantelias, A. & Terzoudi, G. I. Development of an automatable micro-PCC biodosimetry assay for rapid individualized risk assessment in large-scale radiological emergencies. Mutat. Res. Toxicol. Environ. Mutagen. 836, 65–71 (2018).

9. Yadav, U., Bhat, N. N., Shirsaath, K. B., Mungse, U. S. & Sapra, B. K. Refined premature chromosome condensation (G0-PCC) with cryo-preserved mitotic cells for rapid radiation biodosimetry. Sci. Rep. 11, 13498 (2021).

10. Hernansaiz-Ballesteros, R. D., Földi, C., Cardelli, L., Nagy, L. G. & Csikász-Nagy, A. Evolution of opposing regulatory interactions underlies the emergence of eukaryotic cell cycle checkpoints. Sci. Rep. 11, 11122 (2021).

11. Prasanna, P. G. S. & Blakely, W. F. Premature chromosome condensation in human resting peripheral blood lymphocytes for chromosome aberration analysis using specific whole-chromosome DNA hybridization probes. Methods Mol. Biol. 291, 49–57 (2005).

12. Gotoh, E. G2 Premature Chromosome Condensation/Chromosome Aberration Assay: Drug-Induced Premature Chromosome Condensation (PCC) Protocols and Cytogenetic Approaches in Mitotic Chromosome and Interphase Chromatin for Radiation Biology. Methods Mol. Biol. 1984, 47–60 (2019).

13. Gotoh, E. & Durante, M. Chromosome condensation outside of mitosis: Mechanisms and new tools. J. Cell. Physiol. 209, 297–304 (2006).

14. Gelens, L., Qian, J., Bollen, M. & Saurin, A. T. The Importance of Kinase-Phosphatase Integration: Lessons from Mitosis. Trends Cell Biol. 28, 6–21 (2018).

15. Kishimoto, T. Entry into mitosis: a solution to the decades-long enigma of MPF. Chromosoma 124, 417–28 (2015).

16. Hara, M. et al. Greatwall kinase and cyclin B-Cdk1 are both critical constituents of M-phase-promoting factor. Nat. Commun. 3, 1059 (2012).

17. Terakawa, T. et al. The condensin complex is a mechanochemical motor that translocates along DNA. Science (80-.). 358, 672–676 (2017).

18. Hirano, T. Condensins: universal organizers of chromosomes with diverse functions. Genes Dev. 26, 1659–78 (2012).

19. Bazile, F., St-Pierre, J. & D’Amours, D. Three-step model for condensin activation during mitotic chromosome condensation. Cell Cycle 9, 3263–3275 (2010).

20. Kschonsak, M. & Haering, C. H. Shaping mitotic chromosomes: From classical concepts to molecular mechanisms. BioEssays 37, 755–766 (2015).

21. Tsuji, S. & Kanda, R. Chemically induced premature chromosome condensation in short-term cultured human peripheral lymphocytes: applications to biodosimetry. Biotech. Histochem. 82, 29–34 (2007).

22. Sommer, S. et al. The rapid interphase chromosome assay (RICA) implementation: comparison with other PCC methods. Nukleonika 60, 933–941 (2015).

23. Schmidhuber, J. Deep learning in neural networks: An overview. Neural Networks 61, 85– 117 (2015).

24. LeCun, Y., Bengio, Y. & Hinton, G. Deep learning. Nature 521, 436–44 (2015).

25. Krizhevsky, A., Sutskever, I. & Hinton, G. E. ImageNet classification with deep convolutional neural networks. Commun. ACM 60, 84–90 (2017).

26. Vicar, T. et al. DeepFoci: Deep learning-based algorithm for fast automatic analysis of DNA double-strand break ionizing radiation-induced foci. Comput. Struct. Biotechnol. J. 19, 6465–6480 (2021).

27. Shen, X. et al. High-precision automatic identification method for dicentric chromosome images using two-stage convolutional neural network. Sci. Rep. 13, 2124 (2023).

28. Jang, S. et al. Feasibility Study on Automatic Interpretation of Radiation Dose Using Deep Learning Technique for Dicentric Chromosome Assay. Radiat. Res. 195, (2020).

29. Lecun, Y. & Yoshua, B. Convolutional networks for images, speech, and time-series. in The handbook of brain theory and neural networks (MIT Press, 1995).

30. LeCun, Y., Kavukcuoglu, K. & Farabet, C. Convolutional networks and applications in vision. in Proceedings of 2010 IEEE International Symposium on Circuits and Systems 253–256 (IEEE, 2010).

31. Morana, S. J. et al. The Involvement of Protein Phosphatases in the Activation of ICE/CED-3 Protease, Intracellular Acidification, DNA Digestion, and Apoptosis. J. Biol. Chem. 271, 18263–18271 (1996).

32. Kanda, R., Eguchi-Kasai, K. & Hayata, I. Phosphatase inhibitors and premature chromosome condensation in human peripheral lymphocytes at different cell-cycle phases. Somat. Cell Mol. Genet. 25, 1–8 (1999).

33. Prasanna, P. G. ., Escalada, N. D. & Blakely, W. F. Induction of premature chromosome condensation by a phosphatase inhibitor and a protein kinase in unstimulated human peripheral blood lymphocytes: a simple and rapid technique to study chromosome aberrations using specific whole-chromosome DNA hybridizati. Mutat. Res. Toxicol. Environ. Mutagen. 466, 131–141 (2000).

34. Pathak, R. & Prasanna, P. G. S. Premature chromosome condensation in human resting peripheral blood lymphocytes without mitogen stimulation for chromosome aberration analysis using specific whole chromosome DNA hybridization probes. Methods Mol. Biol. 1105, 171–81 (2014).

35. Gao, L., Lu, X., Liu, M.-M., Li, S. & Liu, Q.-J. Transformed Cell Ratio (TCR): A Novel Parameter for Radiation Dose Estimation in Rapid Premature Chromosome Condensation (PCC) Assay Induced by 0–40 Gy Co-60 Gamma Rays. Health Phys. 123, 492–496 (2022).

36. Walev, I. et al. Delivery of proteins into living cells by reversible membrane permeabilization with streptolysin-O. Proc. Natl. Acad. Sci. U. S. A. 98, 3185–90 (2001).

37. Babiychuk, E. B., Monastyrskaya, K., Potez, S. & Draeger, A. Blebbing confers resistance against cell lysis. Cell Death Differ. 18, 80–9 (2011).

38. Teng, K. W. et al. Labeling Proteins Inside Living Cells Using External Fluorophores for Fluorescence Microscopy. Elife 6, (2017).

39. Vigneron, S. et al. Cyclin A-cdk1-Dependent Phosphorylation of Bora Is the Triggering Factor Promoting Mitotic Entry. Dev. Cell 45, 637–650.e7 (2018).

40. St-Pierre, J. et al. Polo Kinase Regulates Mitotic Chromosome Condensation by Hyperactivation of Condensin DNA Supercoiling Activity. Mol. Cell 34, 416–426 (2009).

41. Abe, S. et al. The initial phase of chromosome condensation requires Cdk1-mediated phosphorylation of the CAP-D3 subunit of condensin II. Genes Dev. 25, 863–74 (2011).

42. Gong, D. & Ferrell, J. E. The roles of cyclin A2, B1, and B2 in early and late mitotic events. Mol. Biol. Cell 21, 3149–61 (2010).

43. Xin, G. et al. Aurora B regulates PP1γ-Repo-Man interactions to maintain the chromosome condensation state. J. Biol. Chem. 295, 14780–14788 (2020).

44. Liu, Q. & Ruderman, J. V. Aurora A, mitotic entry, and spindle bipolarity. Proc. Natl. Acad. Sci. 103, 5811–5816 (2006).

45. Wilkins, B. J. et al. A cascade of histone modifications induces chromatin condensation in mitosis. Science 343, 77–80 (2014).

46. Royba, E. et al. Validation of a High-Throughput Dicentric Chromosome Assay Using Complex Radiation Exposures. Radiat. Res. 199, (2022).

47. Gibcus, J. H. et al. A pathway for mitotic chromosome formation. Science (80-.). 359, (2018).

48. Shintomi, K., Takahashi, T. S. & Hirano, T. Reconstitution of mitotic chromatids with a minimum set of purified factors. Nat. Cell Biol. 17, 1014–1023 (2015).

49. Dyson, S., Segura, J., Martínez-García, B., Valdés, A. & Roca, J. Condensin minimizes topoisomerase II-mediated entanglements of DNA in vivo. EMBO J. 40, e105393 (2021).

50. Mochida, S. & Hunt, T. Protein phosphatases and their regulation in the control of mitosis. EMBO Rep. 13, 197–203 (2012).

51. Matsson, P. & Kihlberg, J. How Big Is Too Big for Cell Permeability? J. Med. Chem. 60, 1662–1664 (2017).

52. Güttinger, S., Laurell, E. & Kutay, U. Orchestrating nuclear envelope disassembly and reassembly during mitosis. Nat. Rev. Mol. Cell Biol. 10, 178–91 (2009).

53. Kong, M. et al. Human Condensin I and II Drive Extensive ATP-Dependent Compaction of Nucleosome-Bound DNA. Mol. Cell 79, 99–114.e9 (2020).

54. Cremer, T. & Cremer, M. Chromosome Territories. Cold Spring Harb. Perspect. Biol. 2, a003889–a003889 (2010).

55. Cremer, T. & Cremer, C. Chromosome territories, nuclear architecture and gene regulation in mammalian cells. Nat. Rev. Genet. 2, 292–301 (2001).

56. Pennarun, G., Picotto, J. & Bertrand, P. Close Ties between the Nuclear Envelope and Mammalian Telomeres: Give Me Shelter. Genes (Basel). 14, 775 (2023).

57. Hatzi, V. I. et al. The use of premature chromosome condensation to study in interphase cells the influence of environmental factors on human genetic material. Sci. World J. 6, 1174–1190 (2006).

58. Heng, H. H. Q. et al. Karyotype Heterogeneity and Unclassified Chromosomal Abnormalities. Cytogenet. Genome Res. 139, 144–157 (2013).

59. Kimura, K., Rybenkov, V. V, Crisona, N. J., Hirano, T. & Cozzarelli, N. R. 13S condensin actively reconfigures DNA by introducing global positive writhe: implications for chromosome condensation. Cell 98, 239–48 (1999).

60. Kimura, K. & Hirano, T. ATP-dependent positive supercoiling of DNA by 13S condensin: a biochemical implication for chromosome condensation. Cell 90, 625–34 (1997).

