## Supporting data for "Multiwell-based G0-PCC assay for radiation biodosimetry"

### Supplementary Data.

***Supplementary Figure S1. CNN Architecture for Dose Reconstruction.*** *Analysis starts with convolutional filters that scan the input hPCC image, followed by a non-linear activation to capture complex patterns. A pooling layer then reduces data dimensionality for efficiency. The process concludes with a fully connected neural network that integrates these features to make predictions of the radiation dose.*

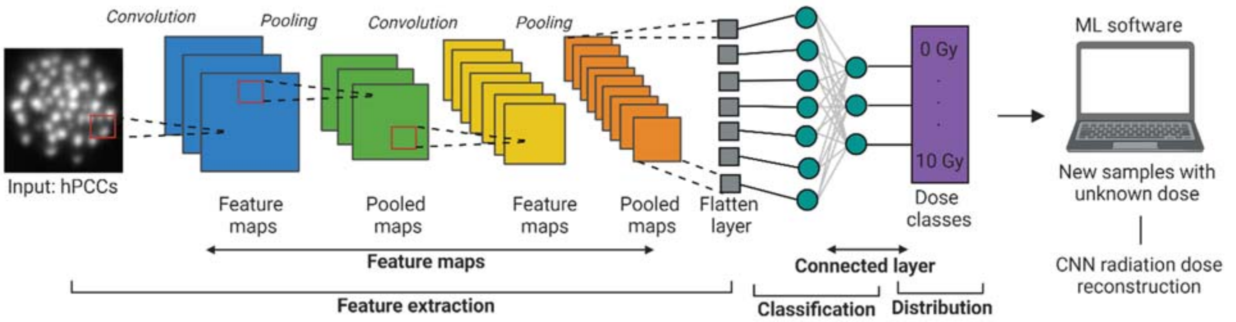


***Supplementary Figure S2. Evaluation of Kinase and SLO Concentrations in the RABiT-II Assay****: (A) Titrating CDK1/Cyclin B kinase with 6.3 nM Cal A. The kinase was diluted starting from 64 µl of kinase per well. The hPCC index was calculated after 6 hours (n = 3). (B) SLO concentration was optimized by titrating it in a 250 µl buffer to achieve 60-80% permeabilization of lymphocytes within 10 minutes. The lymphocyte concentration was maintained between 2.3 to 4.9x10*[6] *cells/mL during the testing. (C) Demonstration of membrane permeabilization with subsequent resealing after 0.2 mg/mL SLO. Propidium Iodide (PI) addition after 10 min. resulted in red fluorescence in permeabilized cells. At 60 min, after membrane repair and Sytox Blue (SB) addition, cells with resealed membranes remained red, while those unable to reseal and undergoing apoptosis fluoresced blue. (D) Fraction of intact, resealed, and dead cells after 0.2 mg/mL and 0.4 mg/mL SLO compared to untreated sample.*

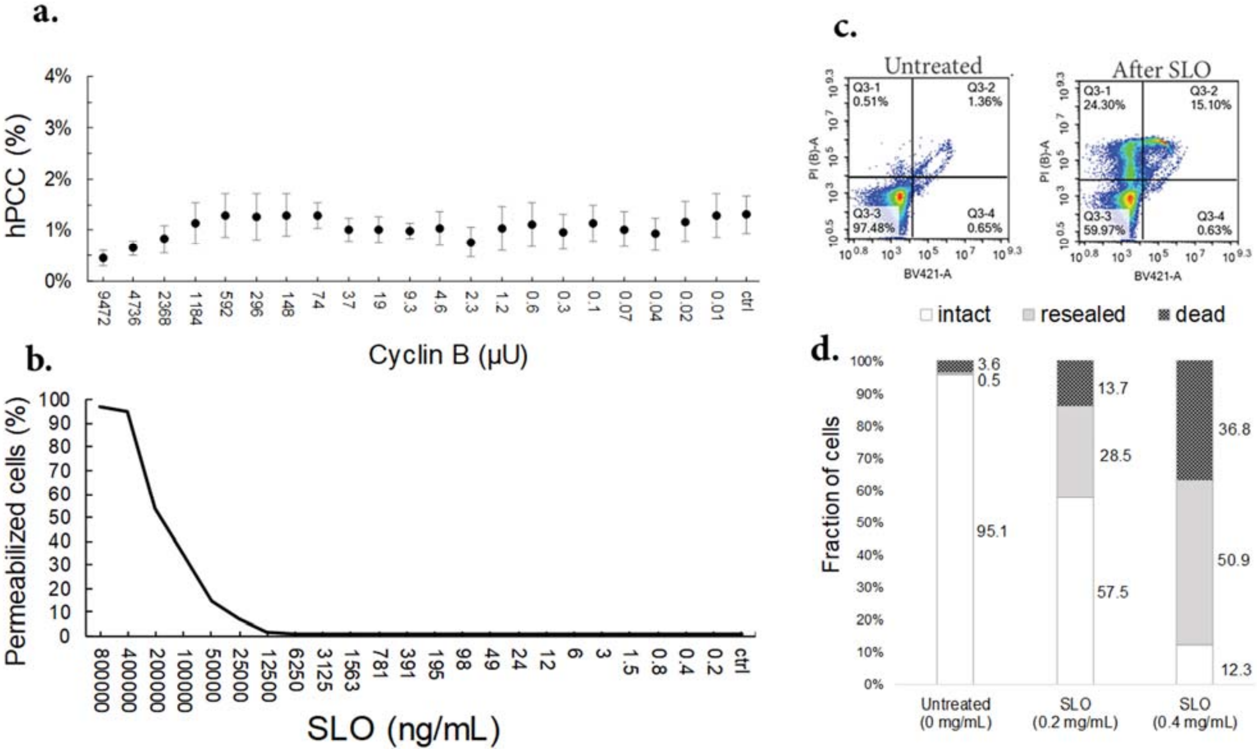

***Supplementary Figure S3.*** *A comparison of chromosome fragmentation in samples exposed to (A) 3 Gy and (B) 8 Gy. On the raw imaged (left side), the representative hPCC are labeled with the letters A, B, C, or D. Enlarged images of these hPCC figures are shown in the inner right inserts. Chromosome pieces within each figure were manually identified and marked with yellow dots. The total count of chromosome pieces for each hPCC is displayed near its corresponding image.*

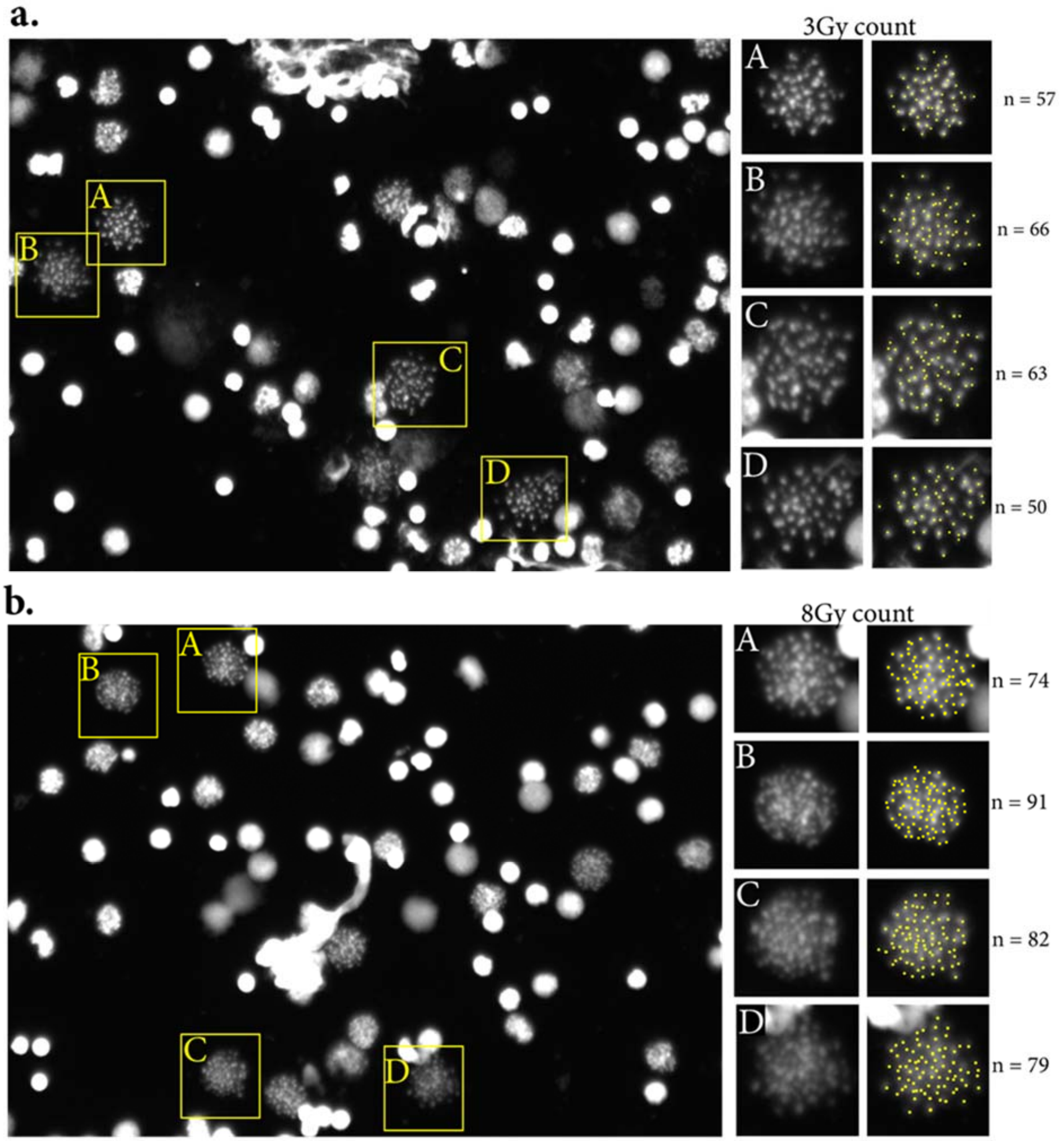

***Supplementary Figure S4.*** *Representative examples of manual scoring of chromosome pieces per hPCC across radiation doses from 0 to 10 Gy. Upper panel: Five random hPCC for each dose (20x, DAPI staining). Bottom panel: Each fragment was manually scored (marked with yellow dots). The resulting number of fragments for each figure is indicated at the bottom of its corresponding image.*

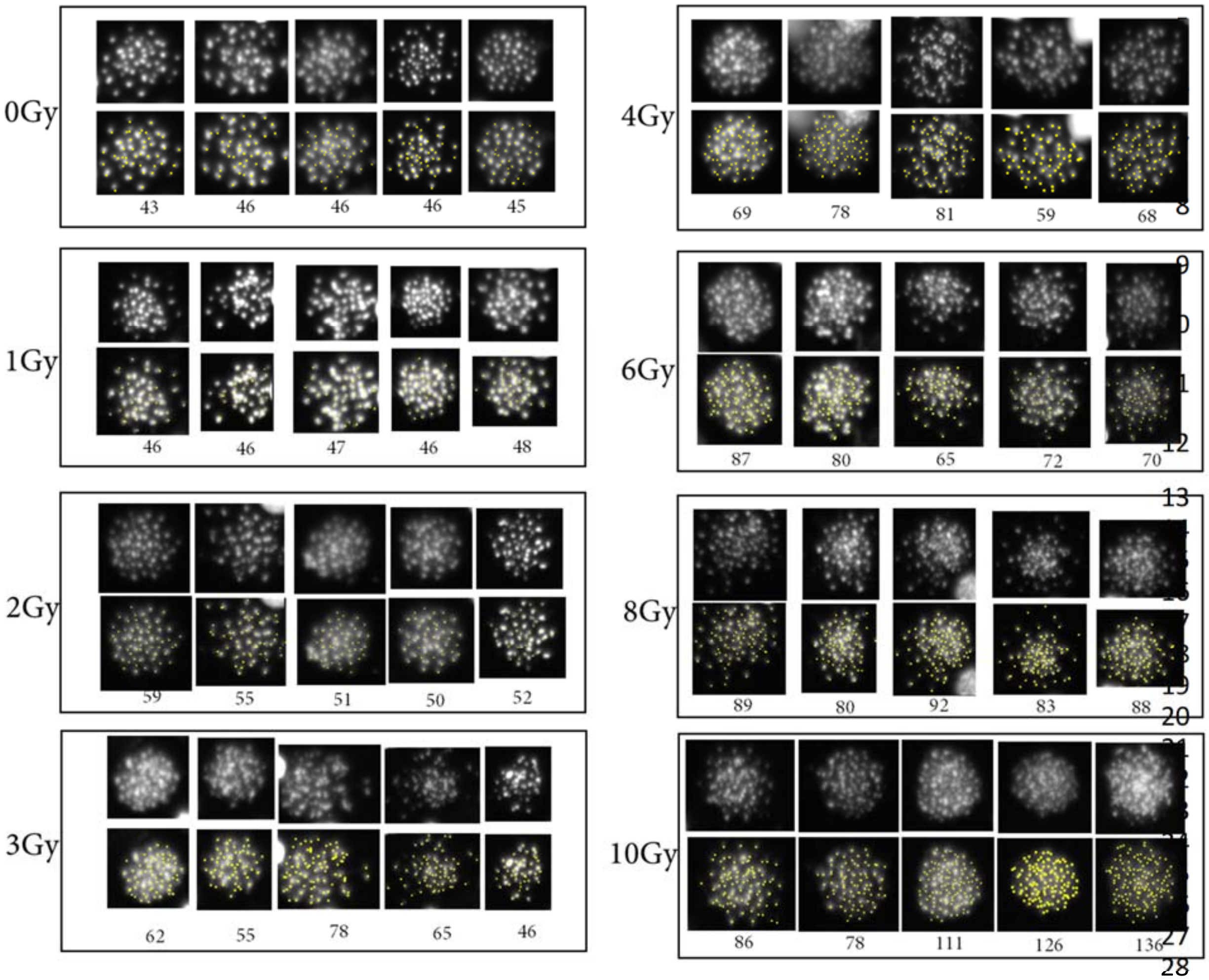


***Supplementary Figure S5.*** *Examples of augmented images generated for CNN training.*

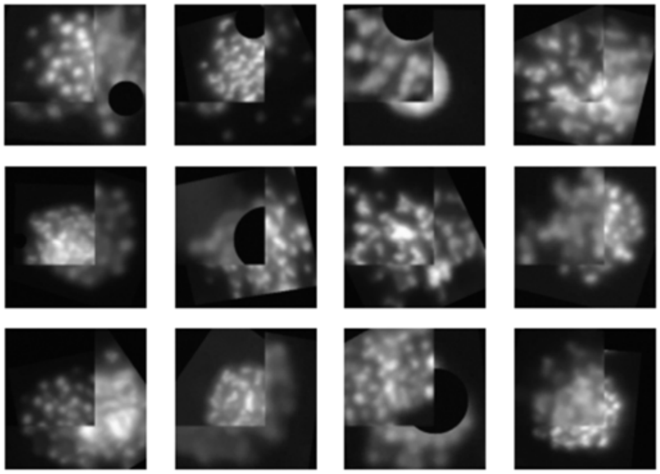

***Supplementary Figure S6. Impact of Cal A Treatment and Radiation on G0-Lymphocytes.*** *The effects of Cal A treatment and radiation on lymphocytes from fresh blood samples of two donors. The lymphocytes were stained with various markers and observed via flow cytometry: DNA damage (H2AX, pSer139), nucleosome state (pHistone H3 Ser10 and pHistone H1.5 Thr154), early apoptosis (PARP Asp214), and cyclin kinase activities (pSer in a CDK substrate motif). (A) Untreated G0-lymphocytes. (B) Treated with 6.3 nM of Cal A. (C) With 50 nM of Cal A. (D) Cells exposed to 8Gy of 320-kVp X-ray, 4Gy/min, without Cal A treatment. (E) Cells are subjected to both, 6.3 nM of Cal A and radiation. (F) 50 nM of Cal A and radiation. Notably, 50 nM of Cal A showed a more pronounced effect on histone phosphorylation compared to 6.3 nM. Both Cal A treatments seem to disrupt DNA repair processes, as indicated by H2AX foci kinetics.*

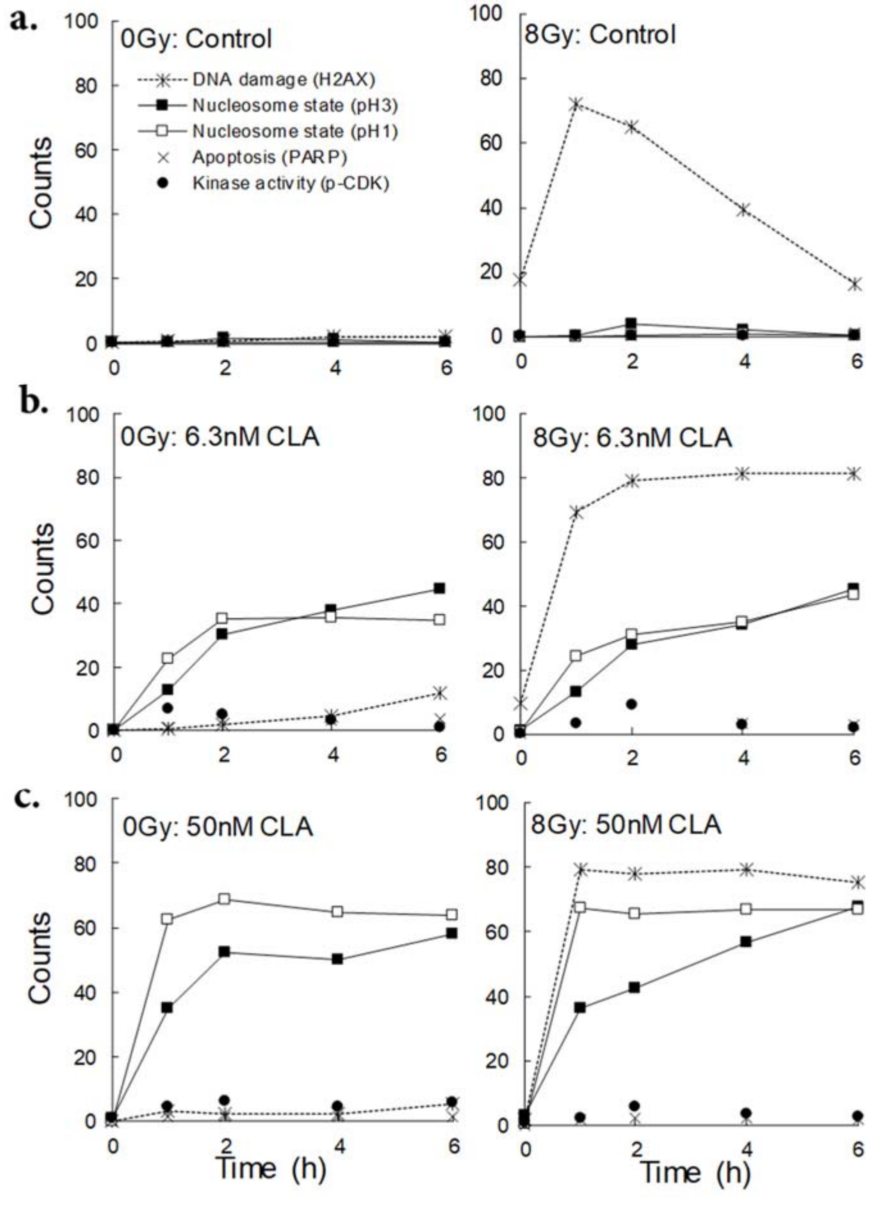

***Supplementary Table S1.*** *Data on chromosome condensation across various Cal A concentrations compile results from 12 independent experiments (10 with 6 hours, and 2 with 7 hours) involving 6 donors (each donor was tested on various days or different durations of Cal A treatment). In each experiment, blood samples were processed in two ways: (i) immediately after collection, or (ii) the following day, after being stored overnight at room temperature. The data for some Cal A concentrations (6400 to 100 nM, and below 3.1 nM) are not included due to the very low or completely absent PCC indices observed at these Cal A concentrations.*

| Donor # (sex/age) | Time (h) | Blood | hPCC index calculation | | | | | | | | | | | | | | |
|---|---|---|---|---|---|---|---|---|---|---|---|---|---|---|---|---|---|
| | | | 50nM Cal A | | | 25nM Cal A | | | 12.5nM Cal A | | | 6.3nM Cal A | | | 3.1nM Cal A | | |
| | | | H | N+L+M | % | H | N+L+M | % | H | N+L+M | % | H | N+L+M | % | H | N+L+M | % |
| # 1 (f, 24) | 6 | fresh | 9 | 19856 | 0.05% | 32 | 14467 | 0.2% | 540 | 78200 | 0.7% | 1206 | 95795 | 1.3% | 425 | 82110 | 0.5% |
| | 6 | o/n | 3 | 15897 | 0.02% | 50 | 42228 | 0.1% | 112 | 66470 | 0.2% | 316 | 78200 | 0.4% | 219 | 62560 | 0.4% |
| # 2 (f, 63) | 6 | fresh | 14 | 34408 | 0.04% | 0 | 0 | 0.0% | 126 | 68816 | 0.18% | 89 | 12512 | 0.71% | 12 | 39100 | 0.03% |
| | 6 | o/n | 152 | 92667 | 0.2% | 329 | 113390 | 0.3% | 556 | 109871 | 0.5% | 692 | 79373 | 0.9% | 96 | 116518 | 0.1% |
| # 3 (f, 30) | 6 | fresh | 36 | 19550 | 0.2% | 37 | 14467 | 0.3% | 185 | 37536 | 0.5% | 409 | 33626 | 1.2% | 209 | 111826 | 0.2% |
| | 6 | o/n | 18 | 20332 | 0.1% | 33 | 31280 | 0.1% | 154 | 43010 | 0.4% | 181 | 23460 | 0.8% | 92 | 50048 | 0.2% |
| # 4 (f, 47) | 6 | fresh | 492 | 367931 | 0.1% | 608 | 49266 | 1.2% | 1638 | 90321 | 1.8% | 2857 | 109089 | 2.6% | 1166 | 93840 | 1.2% |
| | 6 | o/n | 309 | 187680 | 0.2% | 1044 | 89148 | 1.2% | 1896 | 143888 | 1.3% | 1712 | 96968 | 1.8% | 1054 | 111044 | 0.9% |
| | 7 | fresh | 66 | 3128 | 2.1% | 789 | 35190 | 2.2% | 861 | 23460 | 3.7% | 1305 | 31671 | 4.1% | 220 | 9775 | 2.3% |
| # 5 (f, 30) | 6 | fresh | 38 | 26220 | 0.1% | 180 | 43700 | 0.4% | 315 | 69920 | 0.5% | 784 | 58558 | 1.3% | 938 | 113620 | 0.8% |
| | 7 | o/n | 42 | 8740 | 0.5% | 166 | 21850 | 0.8% | 164 | 17480 | 0.9% | 569 | 39330 | 1.4% | 186 | 28405 | 0.7% |
| #6 (f, 65) | 6 | o/n | na | na | na | na | na | na | 229 | 40204 | 0.6% | 298 | 36708 | 0.8% | 485 | 55062 | 0.9% |

Abbreviations: f – female; o/n – overnight; Cal A – Calyculin A; PCC types: N - none, L - low, M - moderate, and H - high condensation; hPCC – index of highly-condensed G0-PCC cells; na – not available.

***Supplementary Table S2.*** *Efficacy of SLO permeabilization. Experiments performed with different SLO badges using fresh blood, on different days, lymphocyte concentration 2.3-4.9x$10^6$ cells/mL (s.e.m., n = 5, 3 donors).*

| SLO (mg/mL) | t = 10 min. | t = 60 min. | | |
|---|---|---|---|---|
| | Permeabilized | Intact | Resealed | Dead |
| | *PI+/SB $_0$* | *PI-/SB-* | *PI+/SB-* | *PI+/SB+* |
| **0.8** | 97 ± 2.8 | 0.5 ± 0.0 | 9.95 ± 6.7 | 89.55 ± 6.7 |
| **0.4** | 94.8 ± 4.2 | 12.28 ± 2.8 | 50.94 ± 2.9 | 36.78 ± 1.8 |
| **0.2** | 54.1 ± 3.1 | 57.48 ± 3.5 | 28.46 ± 2.2 | 13.68 ± 1.7 |
| **0.1** | 34.2 ± 3.2 | 78.8 ± 3.4 | 15.3 ± 2.2 | 5.38 ± 1.4 |
| **0.05** | 14.8 ± 2.7 | 89.72 ± 2.7 | 5.86 ± 2.1 | 3.94 ± 1.1 |
| **0.025** | 7.2 ± 1.8 | 93.92 ± 1.2 | 1.68 ± 0.5 | 4.11 ± 0.9 |
| **0.013** | 1.5 ± 0.8 | 93.77 ± 1.3 | 0.73 ± 0.1 | 4.36 ± 0.9 |
| **0.006** | 1 ± 1.1 | 94.6 ± 1.3 | 0.5 ± 0.0 | 4.3 ± 1.2 |
| **0 (control)** | 0.8 ± 0.2 | 95.06 ± 1.2 | 0.54 ± 0.1 | 3.58 ± 1.0 |

***Supplementary Table S3.*** *Distribution of chromosomal fragmentation among 50 cells per each dose point.*

| #, cell | 0Gy | 1Gy | 2Gy | 3Gy | 4Gy | 6Gy | 8Gy | 10Gy |
|---|---|---|---|---|---|---|---|---|
| **1** | 43 | 46 | 59 | 53 | 69 | 87 | 75 | 86 |
| **2** | 46 | 46 | 50 | 62 | 78 | 80 | 89 | 106 |
| **3** | 46 | 47 | 57 | 55 | 81 | 65 | 80 | 78 |
| **4** | 45 | 46 | 51 | 78 | 59 | 72 | 92 | 111 |
| **5** | 46 | 48 | 60 | 65 | 56 | 75 | 83 | 136 |
| **6** | 45 | 45 | 52 | 46 | 79 | 77 | 75 | 126 |
| **7** | 46 | 46 | 52 | 61 | 71 | 86 | 81 | 107 |
| **8** | 46 | 46 | 50 | 64 | 74 | 73 | 88 | 127 |
| **9** | 46 | 49 | 55 | 76 | 61 | 84 | 80 | 91 |
| **10** | 45 | 47 | 49 | 60 | 75 | 75 | 77 | 92 |
| **11** | 46 | 47 | 58 | 49 | 64 | 71 | 65 | 90 |
| **12** | 46 | 46 | 50 | 58 | 65 | 62 | 78 | 91 |
| **13** | 45 | 52 | 61 | 55 | 65 | 81 | 75 | 79 |
| **14** | 48 | 47 | 59 | 59 | 70 | 60 | 106 | 86 |
| **15** | 46 | 46 | 58 | 53 | 59 | 70 | 76 | 78 |
| **16** | 43 | 52 | 54 | 58 | 57 | 70 | 73 | 79 |
| **17** | 46 | 46 | 60 | 46 | 56 | 61 | 67 | 85 |
| **18** | 46 | 47 | 52 | 70 | 63 | 62 | 70 | 82 |
| **19** | 44 | 44 | 62 | 60 | 59 | 69 | 74 | 87 |
| **20** | 46 | 48 | 50 | 59 | 65 | 78 | 78 | 80 |
| **21** | 43 | 50 | 54 | 52 | 59 | 70 | 79 | 86 |
| **22** | 48 | 50 | 50 | 61 | 61 | 69 | 69 | 88 |
| **23** | 46 | 48 | 54 | 61 | 64 | 82 | 68 | 107 |
| **24** | 46 | 48 | 47 | 67 | 58 | 72 | 83 | 85 |
| **25** | 45 | 46 | 46 | 55 | 60 | 68 | 80 | 90 |
| **26** | 46 | 53 | 59 | 51 | 65 | 67 | 79 | 83 |
| **27** | 47 | 46 | 53 | 46 | 60 | 71 | 73 | 80 |
| **28** | 46 | 46 | 50 | 46 | 59 | 61 | 77 | 93 |
| **29** | 45 | 49 | 50 | 60 | 57 | 80 | 81 | 80 |
| **30** | 46 | 46 | 49 | 57 | 54 | 62 | 75 | 101 |
| **31** | 44 | 46 | 60 | 60 | 73 | 69 | 73 | 87 |
| **32** | 46 | 46 | 50 | 61 | 59 | 52 | 81 | 85 |
| **33** | 46 | 50 | 49 | 54 | 55 | 72 | 85 | 114 |
| **34** | 46 | 53 | 52 | 57 | 59 | 64 | 80 | 109 |
| **35** | 46 | 49 | 58 | 53 | 61 | 69 | 82 | 84 |
| **36** | 46 | 44 | 60 | 50 | 78 | 57 | 83 | 99 |
| **37** | 46 | 49 | 49 | 54 | 50 | 69 | 79 | 88 |
| **38** | 46 | 45 | 57 | 56 | 70 | 67 | 86 | 88 |
| **39** | 43 | 48 | 47 | 57 | 70 | 76 | 77 | 90 |
| **40** | 46 | 46 | 50 | 53 | 53 | 77 | 73 | 98 |
| **41** | 44 | 50 | 54 | 57 | 66 | 77 | 86 | 81 |
| **42** | 46 | 47 | 56 | 66 | 65 | 84 | 77 | 91 |
| **43** | 46 | 47 | 49 | 51 | 63 | 82 | 75 | 97 |
| **44** | 46 | 46 | 52 | 63 | 76 | 76 | 76 | 92 |
| **45** | 45 | 49 | 52 | 59 | 63 | 75 | 82 | 118 |
| **46** | 45 | 48 | 55 | 57 | 64 | 69 | 76 | 73 |
| **47** | 45 | 46 | 59 | 60 | 55 | 68 | 75 | 120 |
| **48** | 46 | 47 | 52 | 52 | 63 | 69 | 80 | 108 |
| **49** | 45 | 48 | 56 | 53 | 61 | 64 | 68 | 115 |
| **50** | 45 | 51 | 47 | 59 | 60 | 59 | 75 | 95 |

### Supplementary Methods

The following sections give information about additional assay optimization experiments and provide a detailed description of our deep learning CNN-based approach for image analysis and dose reconstruction.

##### Reagent Reconstitution

Streptolysin O (SLO) from *Streptococcus pyogenes* (25,000 Units/vial, Sigma Aldrich, MW = 69 kDa) was reconstituted in 1 mL of water to achieve a 1 mg/mL concentration and stored at 2°C for no more than two weeks. Unit is defined as causing 50% lysis of a 2% red blood cell suspension in phosphate-buffered saline at pH 7.4 after 30 minutes at 37°C.

Okadaic acid (OA) from *Prorocentrum concavum* (25 µg, Sigma Aldrich, MW = 805 g/mol) was dissolved in 1 mL of DMSO to reach a concentration of 31 µM and stored at -20°C.

Microcystin-LR (M-LR) solution (10 µg/mL in methanol, Supelco, MW = 995.17 g/mol) was prepared to a final concentration of 10 µM and aliquoted for storage at -20°C.

##### Mitotic Kinases

In our study, we assessed and compared several recombinant human kinases from different suppliers to determine their effectiveness in inducing chromosome condensation for the G0-Premature Chromosome Condensation (G0-PCC) assay. The kinases evaluated include:

From ThermoFisher:

1. CDK1/Cyclin A2 - 10 µg, Molecular Weight (MW): 91 kDa, Catalog #PV6280.
2. CDK1/Cyclin B - 10 µg, MW: 135.2 kDa, Catalog #PV3292.
3. AURKA (Aurora A) - 10 µg, MW: 49.9 kDa, Catalog #PV3612.
4. AURKB (Aurora B) - 10 µg, MW: 43.4 kDa, Catalog #PV6130.
5. PLK1 - 10 µg, MW: 73.3 kDa, Catalog #PV3501.

From Millipore:

6. Cdk1/Cyclin B, Active - 10 µg, MW: 110 kDa, Catalog #14-450.

##### Conversion of Kinase Activity to Units for G0-PCC Assay

Previously, 50 Units of CDK1/Cyclin B from NewEngland BioLabs were found effective in facilitating chromosome condensation in G0-Premature Chromosome Condensation (G0-PCC)[33]. Following the discontinuation of these kinases, we noticed variations in the definition of "Unit" across different kinase vendors, influenced by specific assay conditions detailed in each vendor's Certificate of Analysis. This necessitated a standardization of the enzymatic activity of CDK1/Cyclin B from ThermoFisher, typically quantified in nM/min/mg, into a consistent unit of measurement.

Defining the Unit of Activity: ThermoFisher defines one unit of CDK1/Cyclin B activity as the amount needed to transfer 1,000 nM of phosphate to the Histone H1 peptide substrate PKTPKKAKKL in one minute at 30°C, under the following conditions: 20 mM Tris (pH 7.5), 5 mM MgCl2, 100 µM ATP, 100 µg/mL Histone H1, and trace [32P]-gamma-ATP over a period of 10 minutes in a 50 µL reaction volume.

Calculating Units of Kinase Activity:

1. **Units per mg of Protein:** From the CDK1/Cyclin B analysis, it was determined that 1 mg of this protein transfers 1,480 nmol of phosphate to the Histone H1 substrate. This conversion suggests that 1 mg of protein equates to 1.48 Units (U) of enzyme activity.
2. **Units per Volume:** With the protein concentration at 0.2 mg/mL, each µL contains 0.0002 mg of protein, which corresponds to approximately 0.296 mU/µL. Given a reaction volume of 300 µL in the RABiT-II assay, the addition of 1 µL of this kinase translates to about 0.99 uU/µL.

**Optimization of Streptolysin O Concentration for Lymphocyte Permeabilization**

Optimization of optimal SLO concentration for lymphocyte permeabilization was performed as suggested elsewhere [36]. In brief, to select the concentration of the toxin, 50 µl of cell suspensions (isolated from freshly-drawn blood, and diluted as described in the main Methods section to concentrations 2.3 to 4.9x$10^6$ cells/mL) were resuspended in HBSS without $Ca^{2+}$ containing 30 mM HEPES and serial dilutions of SLO (from 0.8 mg/mL to 0.19 ng/mL) to a final total volume of 250 µl. After 10 min. at 37°C, cells suspensions were mixed with 5 µM of Propidium Iodide (PI) and split into 2 equal parts (125 µl each).

The first aliquout of cells was incubated at room temperature for 10 min. and the percentage of PI-positively stained cells was determined by flow-cytometry (NovoCyte Quanteon). Approximately 30000-40000 events were gated for each titration point. The goal was to determine SLO concentration that causes permeabilization of 60-80% lymphocytes within 10 min of SLO treatment. The second aliquot of cells was subjected to membrane resealing at 37°C, induced by adding $Ca^{2+}$ to the incubation mix. This was done by adding 125 µl of MarrowMax, centrifuging cells for 2 min. at 250 g, washing twice in 200 µl MarrowMax, and final replacing 200 µl with complete MarrowMax culture medium. After 60 min. at 37°C, cells were stained with Sytox Blue (SB), and the percentage of positively stained PI and SB cells was determined by flow-cytometry and gated as described above.

**Kinase Delivery via Reversible Permeabilization**

Based on the results of SLO optimization, lymphocytes were treated with 0.3 mg/mL SLO and presence of various kinases (CDK1/Cyclin B, CDK1/Cyclin A2, Aurora A, Aurora B, and/or PLK1) for 10 minutes, followed by membrane repair and incubation with 6.3 nM Cal A as described in the main Methods section.

**Flow Cytometry**

DNA damage and apoptosis were evaluated during the treatment with Cal A at different time points using H2AX (pSer139) and PARP (cleaved Asp214), both included in the Apoptosis, DNA Damage and Cell Proliferation Kit from BD Pharmingen™ (BD Biosciences, Catalog #562253), following the manufacturer's recommendations. Simultaneously, for analyzing the state of nucleosomes, we used the Phospho-Histone H1.5 (Thr154) Polyclonal Antibody (ThermoFisher) and the Phospho-Histone H3 (Ser10) (D2C8) XP® Rabbit Monoclonal Antibody from Cell Signalling Technologies. In addition, kinase activity was measured using the Phospho-CDK Substrate Motif [(K/H)pSP] MultiMab™ Rabbit Monoclonal Antibody mix (#9477 from Cell Signalling Technologies). Results were analyzed using flow-cytometry (NovoCyte Quanteon) with gated approximately 50000 events per data point.

#### Image Data Pre-Processing

##### *Resizing images*

Resizing of images into 224x224 pixel squares was initiated. The input path, which points to the location of the original images, was defined, along with the output directory where the resized images would be stored. A loop was established to process the images found in the input directory. Each image was read using OpenCV. The calculation of the aspect ratio was performed to determine how to resize the image into a square, while maintaining its original proportions. Depending on the aspect ratio, the image was resized and cropped accordingly. If the resulting image did not meet the specified dimensions, adjustments were made. The current directory was changed to the output directory, and the resized images were saved with new filenames. This process was applied to all the images in the input directory.

##### *Train/test split*

A process to randomly select subsets of images for testing purposes was initiated. A fraction of 30% of the total images was designated for testing, and these images were subsequently moved to a different directory.

##### *Denoising images*

The image denoising process is applied using the 'NonLocalMeansDenoising' class, which is a custom denoising method. The source and destination directories are specified for both training and testing sets. Images from the 'train_dir' and 'test_dir' are used as the source, while the denoised results are saved in 'train_denoised_dir' and 'test_denoised_dir' respectively. The 'NonLocalMeansDenoising' class is responsible for performing the denoising operation. It applies non-local means denoising to the input image, which helps in reducing noise while preserving important details. A function named 'denoise_images' is defined to process the images in the source directory. The denoising process is applied to each image, and the denoised result is saved in the destination directory. The function handles the creation of necessary directories and file paths, ensuring that the denoised images are organized properly. The denoising operation is executed for both the training and testing sets, enhancing the image quality by reducing noise while retaining important details.

#### Image Augmentation

The augmentation code defines two custom classes: 'AddGaussianNoise' and 'BlackoutCircle.' The 'AddGaussianNoise' class is designed to introduce Gaussian noise to an input tensor, while 'BlackoutCircle' generates a blackout circle on an image with a certain probability. These classes serve as custom image transformation techniques, enhancing data augmentation. The 'train_transform' and 'test_transform' transformations are configured for data preprocessing. For training data, transformations like grayscale conversion, random horizontal flips, random affine transformations, and blackout circles with a specific probability are applied. Additionally, Gaussian noise with defined mean and standard deviation is introduced. For testing data, a simpler grayscale conversion and tensor conversion are performed. Training and testing datasets are created using the specified directories and transformations. The 'train_dataset' and 'test_dataset' represent the datasets with applied transformations. Data loaders, 'train_dataloader' and 'test_dataloader,' are established to manage batch processing.

Furthermore, a function named 'cutmix_data' is implemented to apply CutMix augmentation to batches of inputs and labels. This function shuffles inputs and labels, calculates a mixing ratio 'lam,' determines the cropping parameters, and blends the images accordingly. The 'F_Cut_Mix' variable controls the fraction of images to which CutMix augmentation is applied during data iteration. Finally, during the iteration over the training data, CutMix augmentation is employed with a probability determined by 'F_Cut_Mix.' This process involves applying CutMix augmentation to a fraction of the input images in the training dataset, thereby enhancing data diversity and augmenting the training process. In summary, this approach facilitates data augmentation and preprocessing for image datasets, with the capability to apply advanced augmentation techniques like CutMix, Gaussian noise addition, and blackout circles.

For visualization of augmented images, a loop iterates over batches of training data from 'train_dataloader.' Within this loop, the 'cutmix_data' function is applied to introduce CutMix augmentation to 'inputs' and 'labels.' After augmentation, a subset of images is randomly selected for visualization. These selected images are converted from PyTorch tensors to NumPy arrays and have their pixel values rescaled for typical image visualization.

**CNN Model Architecture**

This architecture, encapsulated within the `Net` class, is composed of two fundamental components: a Convolutional Neural Network (CNN) model for feature extraction and a Fully Connected (FC) model for making classification decisions. Within the CNN model, a series of convolutional layers are employed to extract intricate features from the input images. These layers include multiple Conv2d operations, with varying input and output channel dimensions. Notably, the precise configuration of the CNN includes five convolutional layers. Leaky Rectified Linear Unit (Leaky ReLU) activation functions are applied after each convolutional layer, providing the network with non-linearity to capture complex features. Furthermore, max-pooling layers are interleaved after specific convolutional layers, thereby downsizing the spatial dimensions of the feature maps. The CNN model encompasses convolutional layers with filter sizes of 5x5 and 3x3, which are particularly effective for feature extraction at different scales. The convolutional layers are arranged to progressively learn and detect hierarchical features. Within this architecture, some convolutional layers have dropout layers, contributing to better generalization by preventing overfitting.

The Fully Connected (FC) model comprises two linear layers that refine the feature representation and eventually perform classification. The architecture includes Leaky ReLU activation functions in these FC layers, which play a crucial role in the non-linear transformation of feature vectors. Additionally, a dropout layer is introduced in the second linear layer to further regularize the model.

Another noteworthy component is the incorporation of noise augmentation. This process adds controlled random perturbations to the input data, enhancing the model's robustness and its ability to handle variations in input noise. Noise augmentation is an essential aspect of the architecture, promoting the model's adaptability to diverse input conditions. During the forward pass, input data undergoes noise augmentation before being processed through the CNN model. The CNN automatically extracts relevant image features, which are subsequently flattened and reshaped to align with the FC layers. The FC model is responsible for making classification decisions based on the refined feature representation.

The CNN model architecture summary is as follows:

| Layer (type) | Output Shape | | | | Param # |
|---|---|---|---|---|---|
| Conv2d-1 | [-1, | 4, | 220, | 220] | 104 |
| LeakyReLU-2 | [-1, | 4, | 220, | 220] | 0 |
| MaxPool2d-3 | [-1, | 4, | 110, | 110] | 0 |
| Conv2d-4 | [-1, | 8, | 106, | 106] | 808 |
| LeakyReLU-5 | [-1, | 8, | 106, | 106] | 0 |
| MaxPool2d-6 | [-1, | 8, | 53, | 53] | 0 |
| Conv2d-7 | [-1, | 16, | 49, | 49] | 3,216 |
| LeakyReLU-8 | [-1, | 16, | 49, | 49] | 0 |
| MaxPool2d-9 | [-1, | 16, | 24, | 24] | 0 |
| Conv2d-10 | [-1, | 16, | 20, | 20] | 6,416 |
| ELU-11 | [-1, | 16, | 20, | 20] | 0 |
| MaxPool2d-12 | [-1, | 16, | 10, | 10] | 0 |
| Conv2d-13 | [-1, | 32, | 6, | 6] | 12,832 |
| ELU-14 | [-1, | 32, | 6, | 6] | 0 |
| Dropout-15 | [-1, | 32, | 6, | 6] | 0 |
| MaxPool2d-16 | [-1, | 32, | 3, | 3] | 0 |
| Conv2d-17 | [-1, | 64, | 1, | 1] | 18,496 |
| ELU-18 | [-1, | 64, | 1, | 1] | 0 |
| Linear-19 | | | [-1, | 32] | 2,080 |
| LeakyReLU-20 | | | [-1, | 32] | 0 |
| ELU-21 | | | [-1, | 32] | 0 |
| Linear-22 | | | [-1, | 16] | 528 |
| ELU-23 | | | [-1, | 16] | 0 |
| Dropout-24 | | | [-1, | 16] | 0 |
| Linear-25 | | | [-1, | 10] | 170 |

Total params: 44,650
Trainable params: 44,650
Non-trainable params: 0
Input size (MB): 0.19
Forward/backward pass size (MB): 5.66
Params size (MB): 0.17
Estimated Total Size (MB): 6.03

##### Model Training

Class weights are defined to assign different importance to each class in a multi-class classification problem. In this code, a list of class weights is specified, where each weight corresponds to a specific class. These class weights indicate the relative significance of each class during the training process, allowing the model to prioritize or deprioritize certain classes (classes with fewer training images were assigned higher weights to better balance the model's predictions).
